# BCL-X_L_ Dependence is a Subtype Agnostic Actionable Feature of Difficult-to-Treat Kidney Cancers

**DOI:** 10.64898/2025.12.10.693422

**Authors:** Tupa Basuroy, Treg Grubb, Ananthan Sadagopan, Xingping Qin, Kylee Madison, Autumn Hall, Belinda Willard, Sakari Vanharanta, Kristopher A. Sarosiek, Srinivas R. Viswanathan, Abhishek A. Chakraborty

## Abstract

The BCL-X_L_ anti-apoptotic protein is a clear cell Renal Cell Carcinoma (ccRCC) dependency; however, the mechanism of this dependence and its relevance in other aggressive kidney cancer contexts, including metastatic and/or rare RCC subtypes [e.g., Fumarate Hydratase (FH)-deficient and sarcomatoid RCCs], is unknown. Computational predictions, using a machine learning model trained on the human RCC TCGA dataset, and cell-based validations, confirmed BCL-X_L_ dependence in all RCC subtypes. Remarkably, cell state changes, ‘anoikis’ programs, inflammatory state, and metabolic perturbations (e.g., fumarate production in FH-deficient RCCs) independently conferred increased BCL-X_L_ dependence. Correlation studies revealed that increased AMPK isoform 2 (*PRKAA2*) expression is a kidney-specific biomarker of BCL-X_L_ dependence. Indeed, pharmacological AMPK activation sensitized RCCs to BCL-X_L_ blockade. Finally, using functional studies, we developed a multivariate model that accurately predicted BCL-X_L_ dependence in RCC. Our studies offer biomarkers for patient stratification and credential BCL-X_L_ as a subtype agnostic vulnerability in difficult-to-treat RCCs.

## Introduction

Kidney cancer or Renal Cell Carcinoma (RCC) is one of the most common forms of human cancer in both men and women (1). Based on histological and genetic hallmarks, virtually all RCCs can be assigned to one of four subtypes (2, 3): (**a**) clear cell RCCs (ccRCCs), which represent ∼70-75% of all RCCs, and harbor the inactivation of chromosome 3p genes such as von Hippel Lindau (*VHL*), Polybromo 1 (*PBRM1*), and BRCA associated protein 1 (*BAP1*) (4); (**b**) type 1 papillary RCCs (pRCCs), representing ∼5% cases, which show MET activation (5); (**c**) type 2 papillary RCCs, accounting for ∼10% of the tumors, and having a complex set of genetic hallmarks including *PBRM1/BAP1* inactivation or Fumarate Hydratase (*FH*) inactivation; and, (**d**) chromophobe RCC (chRCC), representing ∼5% of the tumors, showing inactivation of Phosphatase and Tensin Homolog (*PTEN*) and Tumor protein 53 (*TP53*). Additionally, the acquisition of sarcomatoid features (a mix of carcinoma and sarcoma-like cells in the same tumor) has been noted in many of these cancer types and is typically associated with worse clinical outcomes (6).

The ccRCCs have been the best studied subtype of kidney cancer because of their clinical frequency and because of the availability of many representative cellular and animal models of human ccRCC. However, the clinical course of the less common RCC subtypes is often more problematic. Despite their varied molecular and histological features, unfortunately, the rarer tumors have been generically categorized into a single bucket as ‘RCCs with variant histology’ or ‘non-ccRCC’. The ‘non-ccRCCs’ typically lack any molecularly defined therapies. Moreover, their clinical management is heavily influenced by guidelines established in ccRCC, which has led to poor clinical outcomes, especially when non-ccRCCs are diagnosed in advanced stages (7). The use of surgery/surveillance is common for localized non-ccRCCs (median 5-year overall survival: ∼ ∼90%), whereas combinations of Tyrosine Kinase Inhibitors (TKIs) and Immune Checkpoint Inhibitors (ICIs) are commonly deployed against advanced disease. Aggressive forms of these cancers, unfortunately, show poor clinical outcomes [e.g., FH-deficient RCC: median Progression Free Survival (PFS) ∼17.2 months (8); sarcomatoid RCC: median PFS ∼6-12 months (9)], highlighting the need for new therapeutics.

The *BCL2L1* gene encodes the anti-apoptotic BCL-X_L_ protein and belongs to a larger family of anti-apoptotic proteins (10), which includes BCL-2 (encoded by the *BCL2* gene), MCL-1 (encoded by the *MCL1* gene), BFL1 (encoded by the *BCL2A1* gene) and BCL-w (encoded by the *BCL2L2* gene) (11). Despite mechanistic similarities in the way that these proteins suppress apoptosis, their expression varies by tissue type, and they regulate cell death in different biological contexts. These proteins also have different binding affinities for pro-apoptotic proteins (e.g., BIM and BID), which further fine-tunes cell death signaling. Interestingly, the role of BCL-X_L_ as a pro-survival mechanism in slow-cycling cells has become evident in recent years. For example, cancer cells experiencing therapy-induced senescence (12–14), cells exiting the cell cycle due to environmental adversity (e.g., nutrient limitation) (15), or cells entering a dormancy/quiescence program via acquisition of stem-like/mesenchymal features (16, 17), have all been independently attributed to show increased BCL-X_L_ dependence.

The oncogenic importance of BCL-X_L_ in kidney cancer has been recently reported in multiple studies. In our previous studies, we discovered that higher BCL-X_L_ dependency was a characteristic of kidney-lineage tumors that display mesenchymal features (17). Similarly, other studies have reported the importance of BCL-X_L_ in metastatic disease (18), and in therapy-induced senescence (13). Unfortunately, all these studies have been limited to ccRCC. Moreover, public databases like DepMap, which, while providing unparalleled access to tumor genetic dependency data (19), currently lack any cells representing the rarer RCC subtypes (e.g., metastatic tumors, FH-deficient RCC, or sarcomatoid RCC). This is a notable knowledge gap, especially considering that these rare RCC subtypes are clinically challenging to control. Studying BCL-X_L_ in other RCC subtypes, such as chRCC, has shown that BCL-X_L_ and its closely related paralog MCL-1 are targetable dependencies in chromophobe RCC (20), lending credibility to the idea that the relevance of BCL-X_L_ blockade outside ccRCC warrants interrogation.

In this study, we began addressing both the mechanisms underlying BCL-X_L_ dependence and the relevance of BCL-X_L_ in rare – but clinically aggressive – forms of kidney cancer. Our studies uncovered that BCL-X_L_ function is critical in many aggressive subtypes of kidney cancer, including metastatic tumors with hyperactivated TGFβ signaling, FH-deficient type 2 pRCCs with supraphysiological fumarate production, and RCCs that are associated with sarcomatoid features. Additionally, using proteomics and transcriptomics approaches, in conjunction with validation studies, we clarified the biological mechanisms of this increased BCL-X_L_ dependence. Relying on public human tumor data (e.g., TCGA) and a machine learning algorithm to predict tumor dependencies based on transcriptome profile of human tumors (21), we next confirmed the relevance of BCL-X_L_ function across human RCC subtypes. Finally, we established and validated subtype-agnostic clinically usable biomarkers of BCL-X_L_ dependence. Altogether, these comprehensive studies offer new mechanistic insights and reinforce the relevance of BCL-X_L_ as a viable target in many difficult-to-treat kidney cancers.

## Results

### Mesenchymal transition rewires the BCL-X_L_ ‘Interactome’

Our previous studies in ccRCC discovered that BCL-X_L_ dependence was increased upon acquisition of mesenchymal features (17); however, the molecular basis of this observation was unclear. We thus began by determining the mechanisms underlying increased BCL-X_L_ dependence in mesenchymal cells. The anti-apoptotic BCL-2 family proteins (e.g., BCL-2, BCL-X_L_, MCL-1, BFL-1 and BCL-w) block cell death by binding the pro-apoptotic BH3-only proteins (e.g., BIM, BID, PUMA, NOXA, etc.) or the pro-apoptotic, pore-forming proteins BAX and BAK that, upon activation, permeabilize the mitochondrial outer membrane to cause release of cytochrome c and initiation of apoptosis (11). Protein-protein interaction (PPI) with the anti-apoptotic BCL-2 family members blocks the function of the pro-apoptotic activator or pore-forming proteins. Therefore, we reasoned that mesenchymal transition reconfigures BCL-X_L_’s interaction with the activator proteins and renders cells vulnerable to BCL-X_L_ blockade.

To address this question, we used the UMRC-2 human ccRCC cell lines. These cells are insensitive to acute exposure to BCL-X_L_ inhibitors (e.g., A1331852); however, exposure to TGFβ, a potent inducer of Epithelial-Mesenchymal Transition (EMT), sensitizes UMRC-2 cells to BCL-X_L_ inhibition (17). We compared drug response curves in these (and following) experiments using a previously published statistical method that deploys a modified Chi-square analysis to compare response curves, independent of their functionality/shape (22). As expected, we noted that TGFβ treatment increased CD44 and decreased CD24 levels (**Fig. 1A**) and correspondingly, in cytotoxicity assays, sensitized these cells to BCL-X_L_ inhibition (e.g., using A1331852) (**Fig. 1B**), but not BCL-2 inhibition (e.g., using ABT199) (**Fig. 1C**).

**Figure 1.**
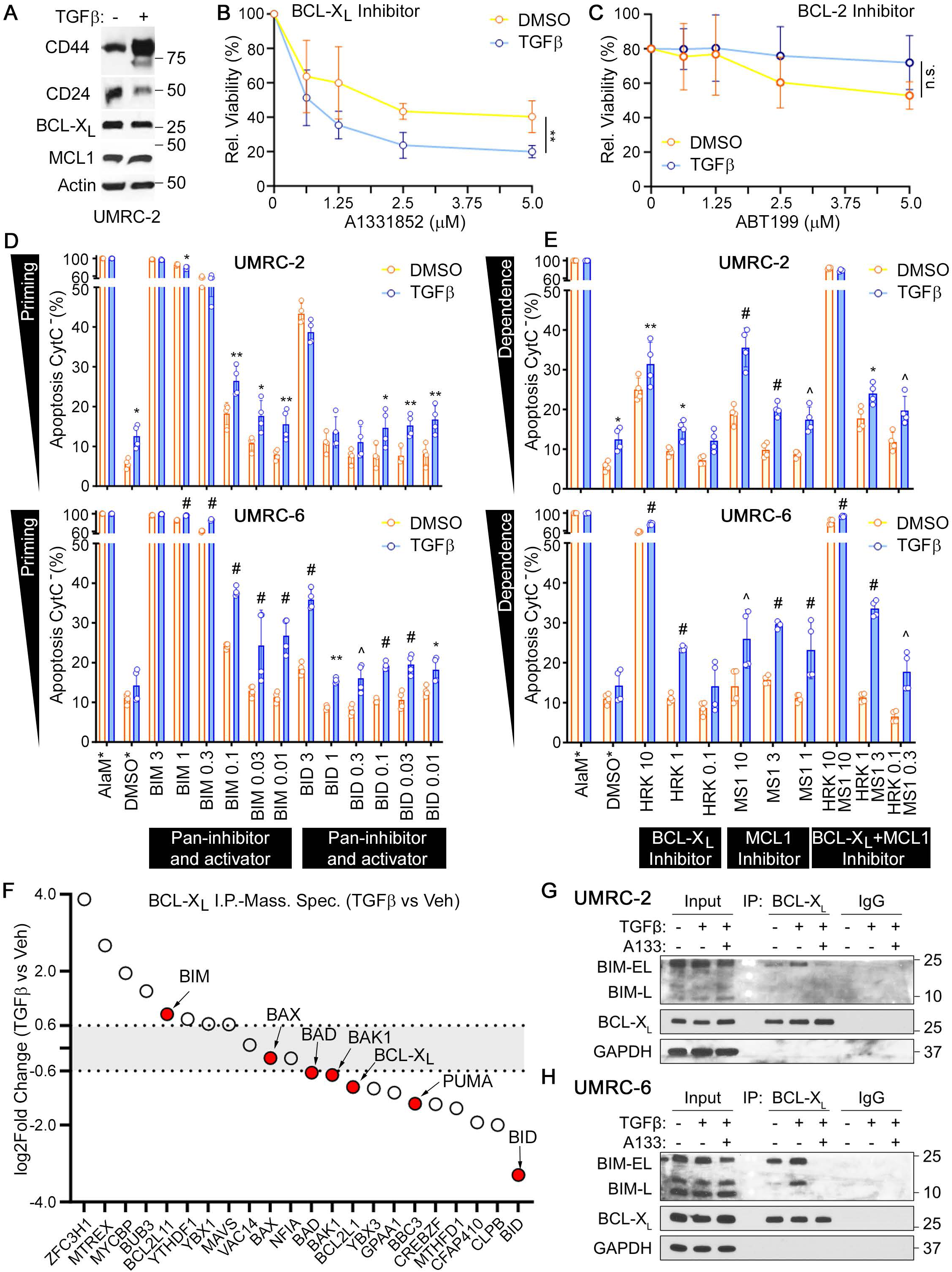
EMT Reprograms the BCL-X_L_ Interactome. (**A**) Immunoblot analysis of UMRC-2 cells that were treated with TGFβ (20 ng/ml) for 4 days. (**B** and **C**) Cell viability measured using CellTiter-Glo following 5-7 days of treatment with the indicated concentrations of the BCL-X_L_ inhibitor, A1331852 (**B**), or the BCL-2 inhibitor, ABT-199 (**C**), for in cells described in (**A**). Curves were compared using the modified Chi-square, curve fit analysis (22); n=3, **p<0.01, n.s.=not-significant. (**D** and **E**) BH3 profiling assays to measure apoptotic response using peptides targeting the activator proteins BIM and BID (**D**) or the anti-apoptotic proteins (HRK peptide: BCL-X_L_ and MS1 peptide: MCL-1), in the indicated ccRCC cell lines that were pre-treated with TGFβ (20 ng/ml) for 4 days. (*) The assays in (**D**) and (**E**) were done in parallel but are presented in two separate panels for clarity. The same positive control (Alamethicin: AlaM) and negative control (DMSO) datapoints are repeated in both the panels for reference. Data was compared using two-way ANOVA, Sidak’s multiple comparisons test; n=3, *p<0.05, **p<0.01, ^p<0.001, #p<0.0001. (**F**) Label-free quantitative mass-spectrometry analysis in cells that were treated with TGFβ (20 ng/ml) for 4 days, or corresponding vehicle control, and BCL-X_L_ binding partners recovered using the anti-BCL-X_L_ antibody or normal IgG (control), as indicated. (**G**) Immunoblot analysis of the indicated proteins in immunoprecipitated complexes recovered using the anti-BCL-X_L_ antibody or normal IgG (control), done using the indicated ccRCC cell lines, that were treated with TGFβ (20 ng/ml) for 4 days ± A1331852 (1 µM) for the last 2 days, where indicated.

We then probed the molecular basis of this change, using the BH3 profiling assays in two human ccRCC cell lines, UMRC-2 and UMRC-6. The BH3 profiling assay measures apoptotic priming and dependencies by detecting the release of cytochrome c from mitochondria after treatment with a panel of pro-apoptotic peptides that disrupt the BH3 domain-dependent PPIs between the anti-apoptotic and pro-apoptotic proteins (23). We found that under control conditions, both the UMRC-2 and UMRC-6 ccRCC cells released cytochrome c in response to moderate (3 μM) or even low (1.0 and 0.3 μM) doses of the pro-apoptotic BIM BH3 peptide (**Fig. 1D**), indicating that these cells were primed for BIM-dependent apoptosis and suggesting that BH3 mimetics are likely to be effective at apoptosis enhancement. Treatment with TGFβ increased cytochrome c release in response to BIM (and, to a lesser degree, BID) BH3 peptides, indicating that this treatment increased apoptotic priming (**Fig. 1D**). Notably, TGFβ treatment also increased the dependence of both these cell lines to BCL-X_L_, as indicated by higher sensitivity to the HRK BH3 peptide, and to a lesser extent increased their dependence on MCL-1, as indicated by higher sensitivity to the MS1 peptide (**Fig. 1E**). Taken together, these BH3 profiling results showed that these apoptotically primed cells became even more primed after TGFβ treatment, becoming especially dependent on BCL-X_L_ for survival.

To determine the cellular changes in BCL-X_L_’s interaction with the activator proteins, we recovered BCL-X_L_ from TGFβ (or vehicle control) treated cells using immunoprecipitation (I.P.) and then performed quantitative (label-free) mass spectrometry analysis. We first processed our dataset to identify proteins that were specifically recovered using the anti-BCL-X_L_ I.P. versus equivalent amounts of the control normal rabbit IgG. We then compared the relative abundance of these proteins in the BCL-X_L_ bound fraction recovered from TGFβ (versus vehicle control) treated cells. These studies demonstrated that TGFβ treatment induced a notable change in the BCL-X_L_ ‘interactome’ and increased BCL-X_L_’s association with BIM (**Fig. 1F** and **Supplementary Table 1**). We then validated these results in both UMRC-2 and UMRC-6 cell lines and confirmed that TGFβ treatment increased the association of BCL-X_L_ with all expressed forms of BIM (e.g., BIM-EL in UMRC-2 cells and BIM-EL/BIM-L in UMRC-6 cells) (**Fig. 1G** and **1H**). Finally, treatment with A133852 disrupted this increased interaction (**Fig. 1G** and **1H**). We thus concluded that TGFβ signaling increases BCL-X_L_’s association with BIM and thereby protects these cells from apoptosis. Disrupting this binding using a BCL-X_L_ targeting BH3 mimetic releases BIM and promotes apoptotic cell death in mesenchymal cells.

### ‘Anoikis’-like programs increase BCL-X_L_ Dependence

EMT is regulated by the concerted action of many mesenchymal transcription factors, including SNAIL and TWIST, and transcriptional reprogramming is prominently associated with this cell state change (24). We thus hypothesized that certain transcriptional programs regulated by TGFβ might render cells vulnerable to BCL-X_L_ inhibition. To this end, we analyzed the alterations in gene expression programs in the UMRC-2 cells and annotated differentially expressed genes (DEGs) that were altered upon TGFβ treatment (≤ ±1.5-fold change, P_adj_ ≤ 0.05) and identified ∼1140 upregulated and ∼540 downregulated genes (**Fig. 2A** and **Supplementary Table 2**). Gene Set Enrichment Analysis (GSEA) (25), using the hallmark gene-sets and the c2 (chemical/genetic perturbation) gene sets, described in the Molecular Signatures Database (mSigDB) (26), showed the expected enrichment of genes associated with TGFβ signaling and EMT, but also hypoxia, perhaps because oxygen deficiency is known to promote EMT (**Fig. 2B** and **Supplementary Table 3**). In contrast, we saw decreased expression of genes in the oxidative phosphorylation and xenobiotic metabolism pathways (**Fig. 2C** and **Supplementary Table 3**).

**Figure 2.**
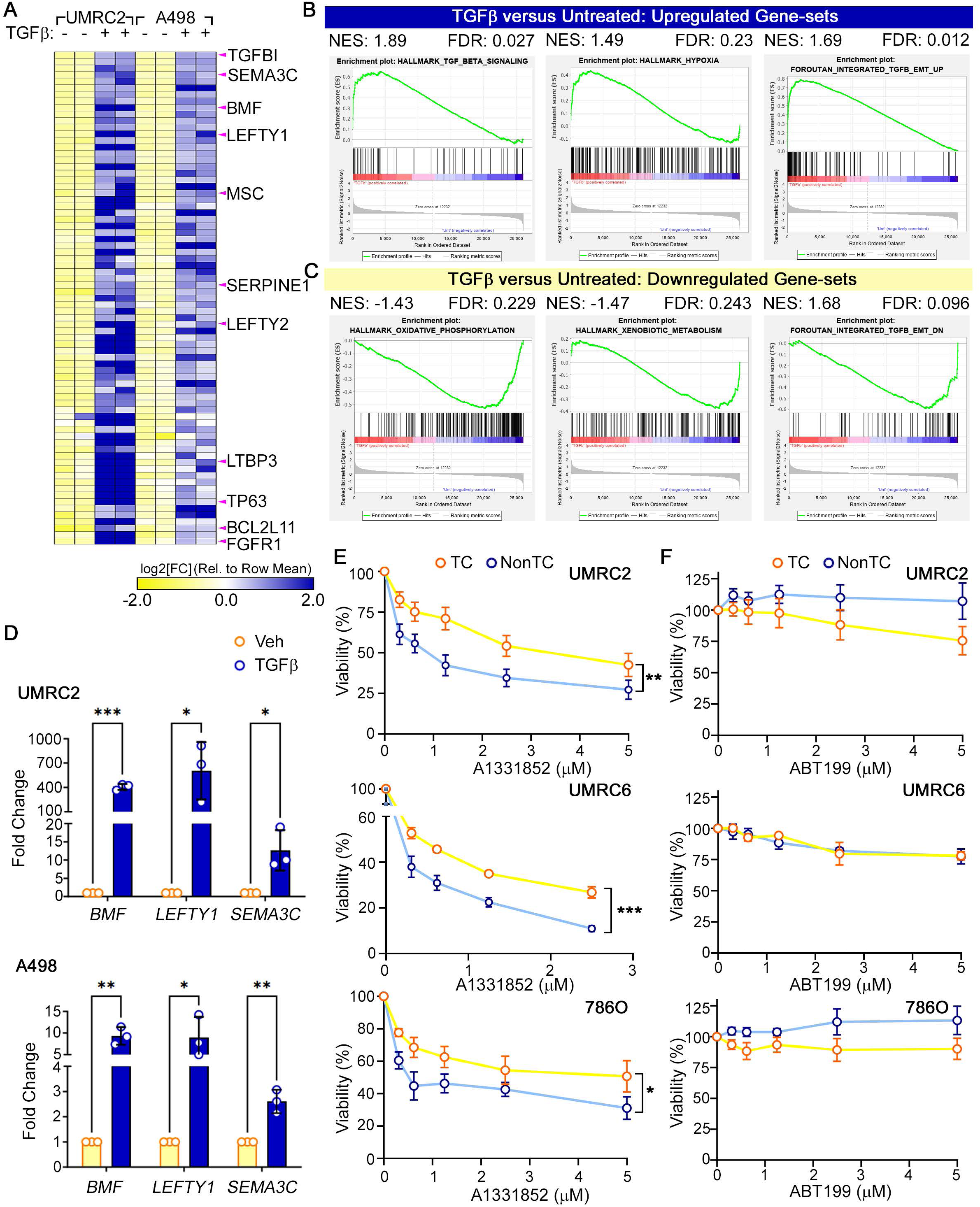
Matrix Detachment Programs Induce BCL-X_L_ Dependence. (**A** to **C**) Heatmap indicating differentially expressed genes (**A**) and GSEA of genes that are upregulated (**B**) or downregulated (**C**) upon treating UMRC-2 and A498 cells with TGFβ (10 ng/ml) for 4 days. (**D**) Fold change in mRNA abundance of cells described in (**A**), as measured using Realtime qPCR analysis, of the indicated genes. Data were first normalized to the abundance of housekeeping genes (β-Actin) and then to the vehicle (control). Data was compared using multiple t-tests (Veh vs TGFβ), corrected using the Holm-Sidak’s method; n=3, *p<0.05, **p<0.01, ***p<0.001. (**E** and **F**) Cell viability measured using XTT following 5-7 days of treatment with the indicated concentrations of the BCL-X_L_ inhibitor, A1331852 (**E**), or the BCL-2 inhibitor, ABT-199 (**F**), in the indicated ccRCC cell lines that were grown on conventional tissue culture (TC) treated plastic or on non-tissue culture (NonTC) plates. Curves were compared using the modified Chi-square, curve fit analysis (22); n=3, *p<0.05, **p<0.01, ***p<0.001.

As a control in these studies, we used the A498 human ccRCC cells. We previously noted that these cells are more mesenchymal than UMRC-2 cells, as evident from their stronger CD44 expression, and correspondingly much more sensitive to BCL-X_L_ blockade (17). Despite these differences in basal mesenchymal state, both cells remain responsive to TGFβ, and low-throughput qPCR studies confirmed that the mesenchymal targets (e.g., *SEMA3C* and *LEFTY1*) showed robust induction upon TGFβ treatment (**Fig. 2D**).

Among the genes most highly induced by TGFβ, we found the BCL-2 modifying factor (*BMF*). BMF controls BCL protein activity and is known to regulate cell death upon activation of ‘anoikis’ (meaning homeless) programs (27) – a quality-control mechanism to trigger cell-death in epithelial cells that are shed from tissue lining. EMT biologically mimics cellular shedding and metastatic cancer cells that are released from epithelial layers need to counteract anoikis-dependent cell death programs.

We hypothesized that BMF induction could regulate TGFβ dependent sensitization to apoptosis via modifications of BCL-X_L_ activity. To address this link, we inactivated BMF using CRISPR/Cas9, then treated these cells with TGFβ to induce EMT, and finally tested their response to the BCL-X_L_ inhibitor. BMF was robustly activated by TGFβ treatment, and its inactivation did not impact EMT programs (**Supplementary Fig. S1A** and **S1B**). However, although we noted some protection in BMF-deficient cells (e.g., BMF sg5 cells lost TGFβ response), inactivation of BMF failed to consistently counteract TGFβ dependent sensitization to BCL-X_L_ blockade (**Supplementary Fig. S1C** to **S1F**). We thus concluded that elevated BMF activity in TGFβ-treated cells was alone not sufficient to regulate BCL-X_L_ response.

We then addressed whether a more global state of ‘homelessness’ sensitizes cells to BCL-X_L_ inhibition. Here, we exploited the fact that seeding adherent cells onto non-tissue culture (NonTC) treated plates enables growth in a semi-detached state that recapitulates anoikis-like features. Using three human ccRCC cell lines that are relatively insensitive to the BCL-X_L_ inhibition (e.g., 786O, UMRC-2, and UMRC-6), we addressed changes in BCL-X_L_ dependence when cells were cultured on traditional tissue culture (TC) dishes versus NonTC dishes and observed a consistent increase in sensitivity to BCL-X_L_ blockade in NonTC conditions (**Fig. 2E**). Importantly, these effects were specific because NonTC conditions neither led to changes in response to the BCL-2 inhibitor, ABT199 (**Fig. 2F**), nor to standard chemotherapeutics (**Supplementary Fig. S2**). We thus concluded that although BMF activitation alone was not sufficient to confer sensitivity to BCL-X_L_ blockade, a more global anoikis-like program did indeed increase BCL-X_L_ dependence in kidney cancer cells.

### Metastatic proficiency promotes sensitivity to BCL-X_L_ blockade

An increased EMT signature is a typical hallmark of metastatic cancer cells. Our observations linking TGFβ dependent EMT and anoikis-like programs with increased sensitivity to BCL-X_L_ inhibition suggested that metastasis-proficiency might also, similarly, promote BCL-X_L_ dependence. The 786M1 cells represent a metastasis-proficient model of the 786O human ccRCC cells (28). Interestingly, although the parental 786O cells are relatively insensitive to BCL-X_L_ inactivation, they were sensitized to BCL-X_L_ blockade on NonTC cells (**Fig. 2E**), suggesting that inducing metastatic proficiency could increase their response to BCL-X_L_ inhibitors.

As expected, the metastasis proficient 786M1 cells showed strong EMT signatures (e.g., increased CD44, TWIST, and Vimentin) (**Fig. 3A**). Remarkably, in cytotoxicity assays, the 786M1 cells showed significantly stronger response to the BCL-X_L_ inhibitor (A1331852) than 786O cells, their non-metastatic counterparts (**Fig. 3B**). These effects were, once again, specific because we did not see any sensitization to the BCL-2 inhibitor, ABT199 (**Supplementary Fig. S3A**).

**Figure 3.**
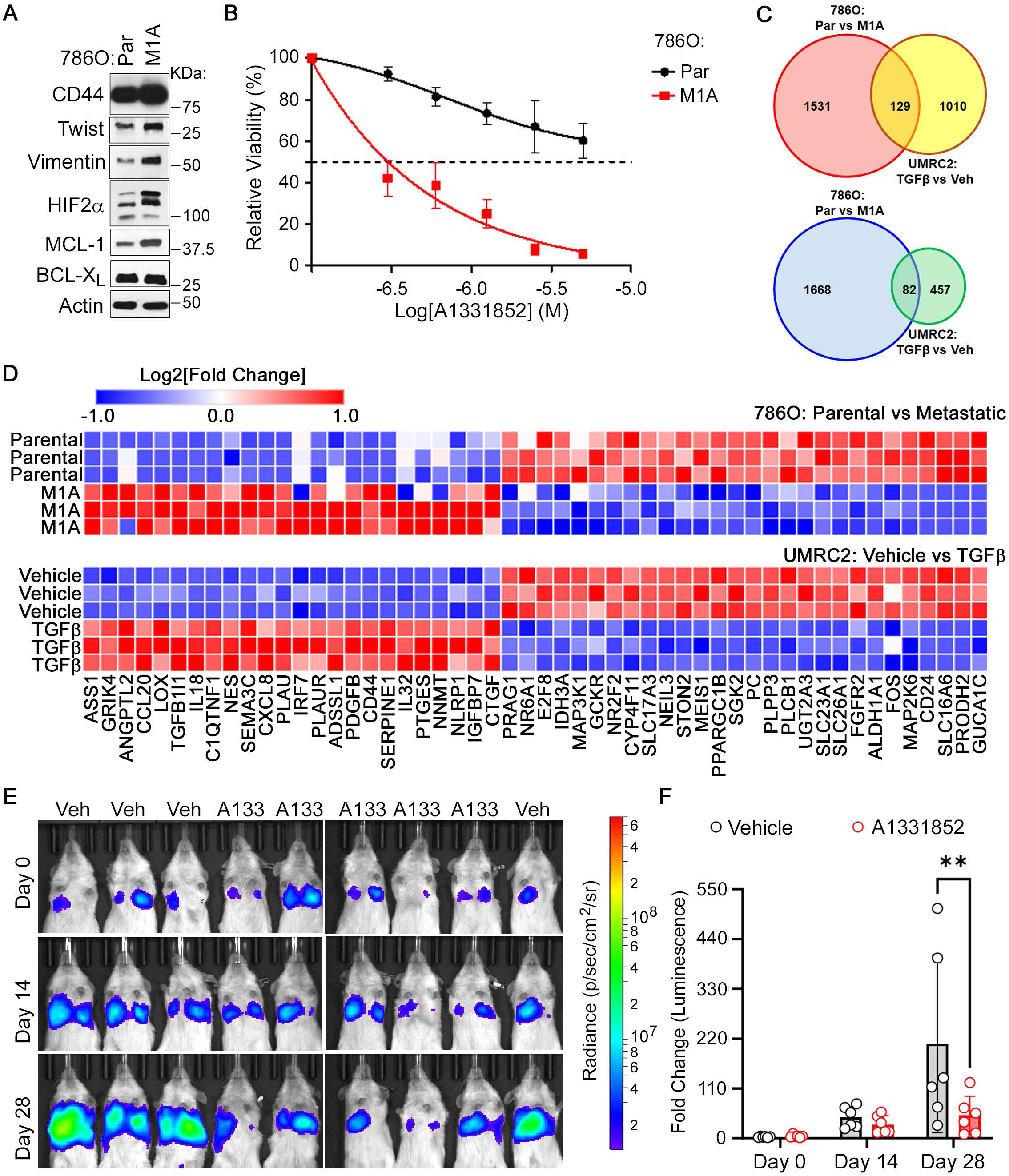
Metastatic Proficiency is Associated with Increased BCL-X_L_ Dependence. (**A** and **B**) Immunoblot analysis of the indicated proteins (**A**) and cell viability (**B**) measured using XTT after 5-7 days of treatment with the indicated concentration of the BCL-X_L_ inhibitor, A1331852, in the metastatic 786M1 cells versus their non-metastatic counterparts, 786O. (**C** and **D**) Venn diagram (**C**) and heatmap (**D**) or genes that are differentially expressed upon TGF treatment in the UMRC-2 cells (as described in Fig. 2A) or in the 786M1 cells (versus 786O). (**E** and **F**) Representative bioluminescence images (**E**) and quantification of bioluminescence signal (**F**) in lung metastatic tumors generated upon tail vein injection of 786M1 cells into NSG mice. Tumors were engrafted for 3-4 weeks following which A1331852 (or vehicle control) was dosed by oral gavage (25 mg/kg, *b.i.d.*) for 4 weeks. Data was compared using two-way ANOVA, Sidak’s multiple comparisons test; Veh (n=5) versus A1331852 (n=6), **p<0.01.

The 786M1 model system offered us a second point of comparison to improve the predictive power of the transcriptional signatures associated with BCL-X_L_ hyper-dependence. We thus generated transcriptomics signatures from the 786O and the 786M1 cells and overlapped the differentially regulated genes in this context with the DEGs identified in the TGFβ treatment experiments. Interestingly, BMF did not feature as an overlapping gene; however, besides the expected higher EMT signature, we found that both the TGFβ treated UMRC-2 cells and the metastatic 786M1 cells showed elevated inflammatory signatures with higher expression of many cytokines, including IL-18 and IL-32 (**Figs. 3C** and **3D**, and **Supplementary Table 4**). Unfortunately, administration of these cytokines individually was not sufficient to confer sensitivity to BCL-X_L_ inhibition in 786O cells (**Supplementary Fig. S3B**). Therefore, we concluded that global changes triggered by TGFβ signaling/metastasis, including an inflammatory state, were necessary to promote BCL-X_L_ dependence.

Tumor metastasis is arguably one of the most common clinical causes of cancer-related mortality. Therefore, we addressed the therapeutic utility of BCL-X_L_ blockade in impeding the growth of metastatic RCCs. To this end, we inoculated the 786M1 cells via the tail vein of immune-deficient mice and followed metastatic tumor engraftment and growth in the lung using bioluminescence imaging. Upon stable tumor growth for ∼20-30 days, we randomized mice to receive A1331852 (25 mg/kg, twice daily) or vehicle control and continued to monitor tumor growth in the lung for an additional 4 weeks. These studies found that BCL-X_L_ blockade led to significantly lower tumor burdens even in the established metastasized lung nodules (**Fig. 3E** and **3F**). Altogether, these studies demonstrated the importance of BCL-X_L_ as an actionable target in clinically aggressive metastatic tumors.

### Inactivation of Fumarate Hydratase (FH) increases response to the BCL-X_L_ inhibitor

All our preceding studies were done in ccRCCs. However, we recently reported that targeting BCL-X_L_, in concert with MCL-1 blockade, was also an apoptotic vulnerability of certain chRCCs (20). Additionally, the clinical control of rare (non-ccRCC) subtypes of kidney cancer remains a significant unmet need (7). We thus addressed the importance of BCL-X_L_ dependence in other rare RCC subtypes.

We first focused on the Fumarate Hydratase (FH)-deficient hereditary leiomyomatosis RCCs (HLRCCs), which are classified as type 2 pRCCs. These tumors often occur in young patients, lack any known custom therapeutic vulnerabilities, and remain challenging to combat in the clinic. FH canonically functions in the tricarboxylic acid (TCA) cycle where it catalyzes the conversion of fumarate into malate in the mitochondria. FH inactivation, therefore, causes profound mitochondrial dysfunction, metabolic rewiring, and cellular perturbations due to the hyperaccumulation of fumarate (29). Fumarate accumulation in FH-deficient tumors has been reported to drive elevated mesenchymal signatures (30). Moreover, mitochondrial dysfunction is often associated with altered dependence on apoptotic programs. Therefore, we addressed if BCL-X_L_ dependence was a feature of these tumors.

First, we genetically eliminated FH activity in the TK-10 human non-ccRCC cells that have pRCC features (31). We lentivirally transduced these cells with CRISPR/Cas9 constructs to target FH (sgFH) or two non-targeting controls (sgNT1 and sgNT2). After selection in Puromycin for 7-10 days, we confirmed efficient elimination of FH by western blotting (**Fig. 4A**) and then seeded these cells to measure response to BCL-X_L_ and BCL-2 inhibitors. These studies confirmed that FH inactivation conferred sensitivity to BCL-X_L_, but not BCL-2, inhibition (**Fig. 4B** and **Supplementary Fig. S4A**).

**Figure 4.**
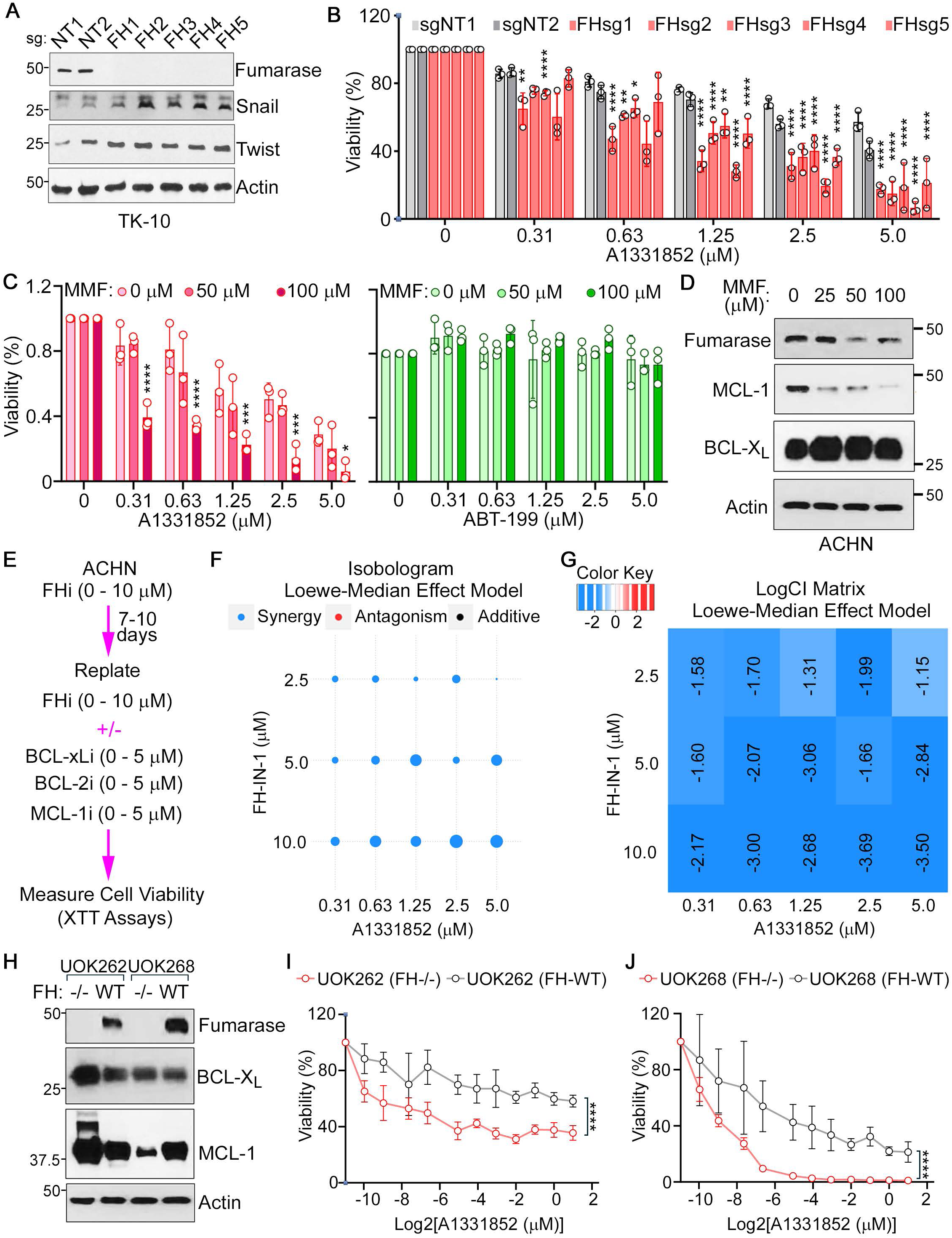
Fumarate Hydratase (FH) Inactivation Sensitizes RCCs to BCL-X_L_ Blockade. (**A** and **B**) Immunoblot analysis (**A**) and cell viability assays (**B**), done using CellTiter-Glo, in TK-10 cells that were lentivirally transduced to express sgRNAs targeting FH or non-targeting (NT) controls. In (**B**), cells were selected in Puromycin (2 µg/ml) for 7 days following which they were seeded and treated with the indicated doses of A1331852 for 5-7 days. Data was compared using two-way ANOVA, Dunnett’s multiple comparisons test; n=3, *p<0.05, **p<0.01, ****p<0.0001. (**C** and **D**) Cell viability, measured using XTT assays (**C**), and immunoblot analysis (**D**) of ACHN cells that were pre-treated with the indicated concentrations of Mono-methyl fumarate (MMF) for 14 days and then cultured in the presence of the indicated concentrations of A1331852 or ABT-199 for 5-7 days. In (**C**), data was compared using two-way ANOVA, Dunnett’s multiple comparisons test; n=3, *p<0.05, ***p<0.001, ****p<0.0001. (**D**) was done using cell pellets harvested at the end of the MMF pre-treatment. (**E** to **G**) Schema of experiment (**E**), isobologram (**F**), and LogCI matrix (**G**), plotted using the SiCoDEA algorithm (33), of ACHN cells that were treated with the indicated doses of the FH inhibitor (FH-IN-1) in combination with the indicated doses of A1331852. (**H** to **J**) Immunoblot analysis (**H**) and cell viability measurements, done using CellTiter-Glo, after treatment with the indicated concentrations of A1331852 for 5-7 days, using the FH-deficient UOK262 (**I**) or UOK268 (**J**) that were modified to stably express wild-type FH. Curves were compared using the modified Chi-square, curve fit analysis (22); n=3, ****p<0.0001.

Inactivation of FH leads to an increase in intracellular (and secreted) fumarate levels. Thus, the metabolic effects of FH inactivation can be modeled by subjecting FH-proficient cells to increasing fumarate levels. To this end, we cultured a second human pRCC cell line, ACHN, with two concentrations (50 µM and 100 µM) of a cell-permeable (esterified) version of fumarate [mono-methyl fumarate (MMF)] for 14 days. We then tested the effect of the BCL-X_L_ and BCL-2 inhibitors. Although the lower dose did not elicit significant change in response, we found that 100 µM MMF pre-treatment – which was lower than the typical 200-400 μM levels used in previous studies (30) – was sufficient to sensitize ACHN cells to BCL-X_L_ inhibition, but not BCL-2 inhibition (**Figs. 4C** and **4D**).

Next, we deployed a pharmacological approach to measure the functional links between FH activity and BCL-X_L_ dependence. We first treated ACHN cells with the FH inhibitor, FH-IN-1 (32), for 7-10 days. These pre-treated cells were replated and then continued to be exposed to FH-IN-1 in the presence of either a BCL-X_L_ inhibitor (BCL-X_L_i: A1331852), a BCL-2 inhibitor (BCL-2i: ABT199), or an MCL-1 inhibitor (MCL1i: S63845). We analyzed the effects of these inhibitors using the ‘Single and Combined Drug Effect Analysis (SiCoDEA)’ algorithm (33). These drug-drug interaction studies showed robust synergistic effects between the FH inhibitor and the BCL-X_L_ inhibitor, but not the BCL-2 or MCL-1 inhibitors (**Figs. 4E** to **4G** and **Supplementary Figs. S4B** to **S4C**). If anything, we noted antagonism with FH-IN-1 at several combinations of the BCL-2i and MCL1i.

Two cell lines models, UOK262 (UOK262^FH-null^) and UOK268 (UOK268^FH-null^), are routinely employed as faithful representatives of human FH-deficient HLRCCs (34, 35). Additionally, isogenic FH-proficient stable cell lines (UOK262^FH-WT^ and UOK268^FH-WT^), with physiological levels of FH expression restored, have been previously generated. We deployed both these models in our studies. After confirming expected expression patterns of FH (also called Fumarase) using western blotting (**Fig. 4H**), we tested BCL-X_L_ dependence. We observed that the FH-null cell lines showed significantly higher sensitivity to the BCL-X_L_ inhibitor than their FH-WT counterparts (**Figs. 4I** and **4J**). Neither the FH-null cells nor their WT counterparts showed any response to the BCL-2 inhibitor (**Supplementary Fig. S4D** and **S4E**). These rigorous studies confirmed that FH inactivation was sufficient to confer sensitivity to BCL-X_L_ blockade, thus justifying the use of BCL-X_L_ agents against these (often) aggressive renal cancers.

### Predictive Value of DepMap Correlations on BCL-X_L_ dependence

The Cancer DepMap is primarily represented by common tumor subtypes – and in the case of kidney cancer, predominantly ccRCC cells. To address this gap, we leveraged a recently developed machine learning model ‘Transcriptional Prediction of Lethality (TrPLet)’ that predicts gene dependency scores based on tumor or cell line transcriptome data (21). Applying TrPLet to predict the importance of the BCL family proteins in RCC subtypes, using RCC RNA-Seq data from the TCGA, we found that BCL-X_L_ (encoded by *BCL2L1*) was predicted to be a notable dependency across all RCC subtypes. In contrast, genes that encoded BCL-2 (*BCL2*), MCL-1 (*MCL1*), or BCL-w (*BCL2L2*), showed no predicted dependence in kidney tumor cells versus other lineages (**Fig. 5A**).

**Figure 5.**
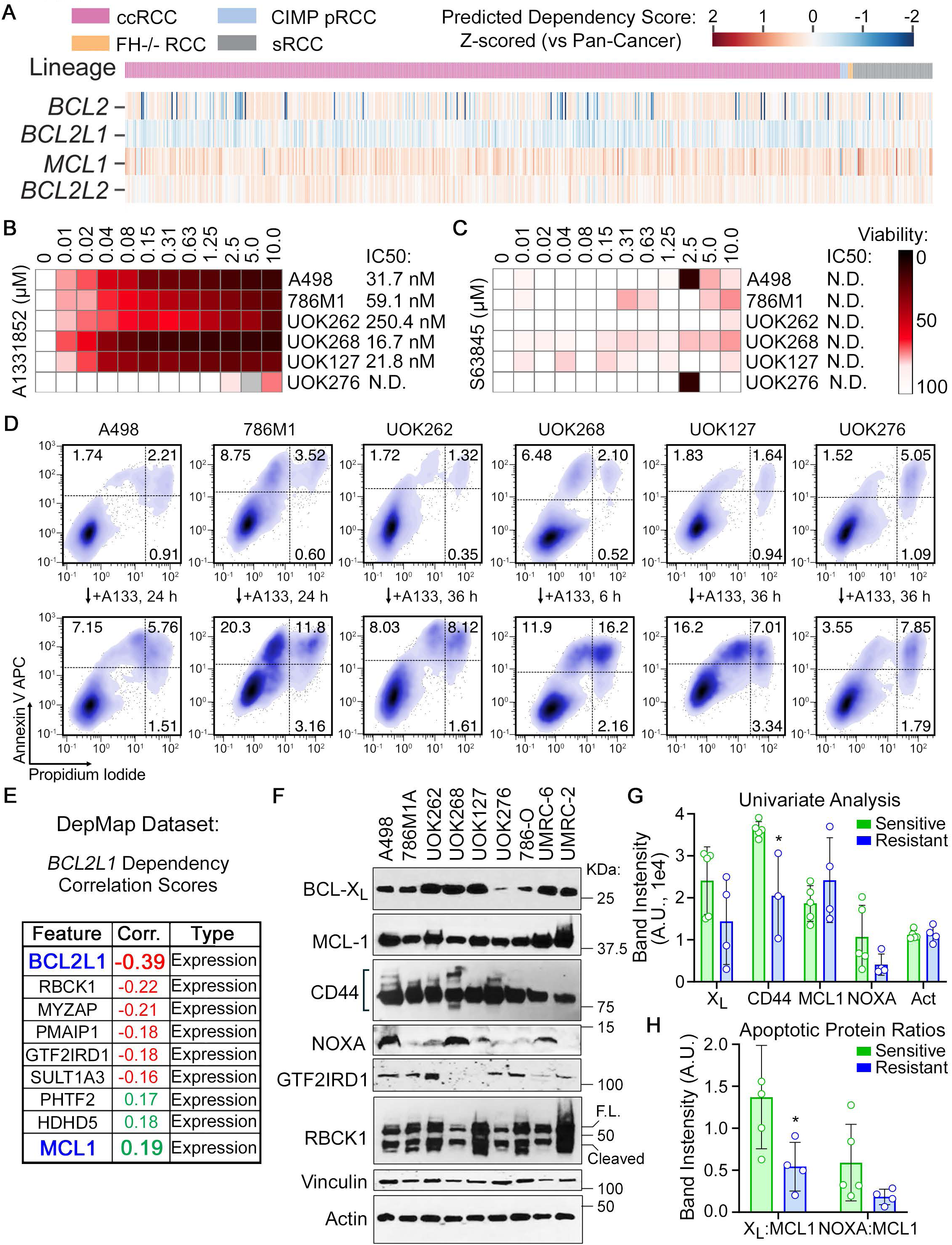
Computational Tools Establish the Relevance of BCL-X_L_ Dependency in Human RCCs and Offer Correlative Biomarkers. (**A**) Machine learning analysis, using the TrPLet algorithm (21), of transcriptomics signatures in the TCGA renal cancer dataset to identify the relative dependencies in the BCL-2 family proteins. (**B** and **C**) Heatmaps representing cell viability of the indicated RCC cell lines that were treated with the BCL-XL inhibitor (A1331852) (**B**) or the MCL-1 inhibitor (S63845) (**C**) for 7 days. IC50 measurements were made using regression analysis in GraphPad Prism. (**D**) Flow cytometry analysis (AnnexinV vs PI) of the indicated cells that were treated with 100 nM A1331852 (A133) for the indicated time. (E) Predictability scores derived from the DepMap dataset for genes that correlate with higher BCL-XL dependence (more negative Chronos scores). (F to H) Immunoblot analysis (F), univariate analysis (G), and apoptotic protein ratios (H), done using ImageJ based quantifications of the indicated proteins in the renal cancer panel. Data was compared using two-way ANOVA, Sidak’s multiple comparisons test; BCL-X_L_i sensitive (green, n=5) versus resistant (blue, n=4), *p<0.05.

We then validated these computational predictions. We deployed a BCL-X_L_ dependent non-metastatic ccRCC (A498), a metastatic ccRCC (786M1), and the FH-deficient RCCs (UOK262^FH-null^ and UOK268^FH-null^). Additionally, we expanded these studies to include another category of aggressive RCCs – the sarcomatoid tumors. These tumors are a histological categorization of kidney tumors and can occur in tumors of the other subtypes, which means that these tumors bear very disparate genetic hallmarks. Importantly, acquisition of mesenchymal features is a common feature of sarcomatoid RCCs (36). We thus deployed two models of sarcomatoid RCCs: (**i**) the UOK127 cells, which represent sarcomatoid ccRCCs because they show *VHL* inactivation via promoter hypermethylation but have WT p53 activity (37); and (**ii**) the UOK276 cells, which represent sarcomatoid chRCCs and are *VHL*-proficient but lack p53 activity (38).

We annotated these cells, using pharmacological inhibitors, with respect to their BCL-X_L_, BCL-2, and MCL-1 dependence. These studies showed that, other than the sarcomatoid chRCC (UOK276) cells, all the other cell lines were exquisitely sensitive to BCL-X_L_ loss, showing IC_50_ in the nM range for A1331852 (**Fig. 5B** and **Supplementary Fig. S5A**) but not the MCL1 inhibitor (**Fig. 5C**) or the BCL-2 inhibitor (**Supplementary Fig. S5A**). Moreover, the fitness defects conferred by BCL-X_L_ inhibition occurred via the canonical apoptotic pathway. Apoptosis flips phosphatidylserine (PS) from the inner to the outer side of the membrane, which can be detected using Annexin V (AnnV) staining in non-permeabilized cells. We noted an increase in both the early apoptotic [AnnV^high^; Propidium Iodide (PI)^low^] and the late apoptotic (AnnV^high^; PI^high^) populations in the RCCs treated with the BCL-X_L_ inhibitor (**Fig. 5D**). Here too, the UOK276 cells showed minimal changes in the apoptotic population compared to the other cells in the analysis. These results validated the predictions made using TrPLet and demonstrated the relevance and specificity of BCL-X_L_ dependence across a spectrum of RCC subtypes.

We then probed for molecular biomarkers of BCL-X_L_ dependence in the DepMap database and found that BCL-X_L_ dependence correlated with higher expression of *BCL2L1* (which encodes BCL-X_L_), *GTF2IRD1*, *RBCK1*, and *PMAIP1* (also called NOXA) **Fig. 5E** and **Supplementary Fig. S5B**). In contrast, higher BCL-X_L_ dependence correlated with lower *MCL1* expression (**Supplementary Table 5**). These predictions were based on computational – lineage-agnostic – analysis of BCL-X_L_ dependence across the entire panel of cell lines used in the DepMap project. We thus addressed how these biomarkers scored in kidney cancer cells. Additionally, given our own findings linking mesenchymal state and BCL-X_L_ dependence, we also analyzed how CD44 expression might correlate with BCL-X_L_ dependence (more negative chronos scores). Although the trends for *BCL2L1*, *MCL1*, and *PMAIP1* expression (versus BCL-X_L_ dependence) in kidney cancer cells (red trend lines) resembled the lineage agnostic patterns (gray trend lines) (**Supplementary Figs. S5C** to **S5E**), *CD44* was a notable exception. We found that *CD44* expression had no predictive value when used against the entire DepMap dataset (**Supplementary Fig. S5F**, gray line, r=-0.010); however, in kidney cancer cells, high CD44 expression correlated highly with BCL-X_L_ dependence (**Supplementary Fig. S5F**, red line, r=-0.421).

Identification of relevant patient cohorts using predictive biomarkers can prove invaluable in improving treatment outcomes. Based on our functional annotations, we next addressed the relevance of the DepMap predicted correlations as biomarkers of BCL-X_L_ dependence (**Fig. 5E**) in our RCC panel. To address this question, using western blotting, we measured the protein expression levels of the biomarkers identified in the DepMap analysis (e.g., BCL-X_L_, MCL1, NOXA, etc.). We quantified the protein expression using ImageJ and compared the expression of the individual proteins in cells that were annotated as BCL-X_L_ sensitive (e.g., A498, 786M1, UOK262, UOK268, and UOK127) or BCL-X_L_ insensitive (e.g., UOK276, 786O, UMRC-2, and UMRC-6) (**Figs. 5B** and **5D**). We then performed univariate analysis to determine the predictive power of each individual biomarker. Considering the known mesenchymal properties of sarcomatoid cells, we excluded the CD44 datapoint from the UOK276 cells in this analysis, because this cell line’s p53-deficient status, low BCL-X_L_ expression, and lack of response to BCL-X_L_ blockade created an experimental confounder. This analysis demonstrated that CD44 expression was indeed a significant predictor of BCL-X_L_ dependence in kidney lineage cancer cells (**Fig. 5G**).

Although BCL-X_L_, PMAIP1/NOXA, and MCL-1 expression trended in the direction predicted by the DepMap dataset, these markers individually were unable to demarcate BCL-X_L_ dependent vs independent cell lines (**Fig. 5G**). Therefore, we considered whether co-occurrence of these features might improve our ability to predict BCL-X_L_ dependence. Indeed, higher BCL-X_L_:MCL-1 ratios were seen in the BCL-X_L_ sensitive (versus insensitive) cohorts (**Fig. 5H**). In summary, computational predictions and our validation studies offered potential biomarkers to identify BCL-X_L_ dependent patient cohorts across all RCC subtypes.

### Multivariate Analysis Identifies a Reliable Biomarker Panel of BCL-X_L_ Dependence

The DepMap predicted correlates of BCL-X_L_ dependence showed many genes having a different correlation coefficient in the kidney lineage cells versus the lineage agnostic analysis. Most notably, CD44, which failed to score as a correlate of BCL-X_L_ dependence in the lineage agnostic analysis, had very strong predictive power in the kidney lineage cells. We thus reanalyzed the DepMap data in a lineage-specific manner, using two BCL-X_L_ dependent lineage cells – kidney and bowel cancer. This analysis identified ∼750 genes as correlates of BCL-X_L_ dependence specifically in kidney cancer. We then disregarded hits in the HIF pathway, kidney development, scaffolding proteins, and cytokines because these were likely to be biological artifacts of the chronic HIF/Hypoxia pathway activation or the inflammatory state in mesenchymal kidney tumors. Truncating the gene list in this manner offered ∼400 genes with a Pearson coefficient <-0.35. In this list, we found *SLC1A1*, which we have recently reported as a RCC oncogene associated with elevated expression in metastatic tumors (39). Additionally, we noted *PRKAA2* among the genes whose expression correlated with BCL-X_L_ dependence (**Fig. 6A** and **Supplementary Table 6**). This was specific because, in contrast to RCC, *PRKAA2* failed to score as a correlate of BCL-X_L_ dependence in the bowel or lineage-agnostic analysis (**Supplementary Fig. S6A** and **S6B,** and **Supplementary Table 6**). Moreover, expression of *PRKAA1*, which encodes AMPKα1, did not correlate with BCL-X_L_ dependence (**Figs. 6B** and **6C**).

**Figure 6.**
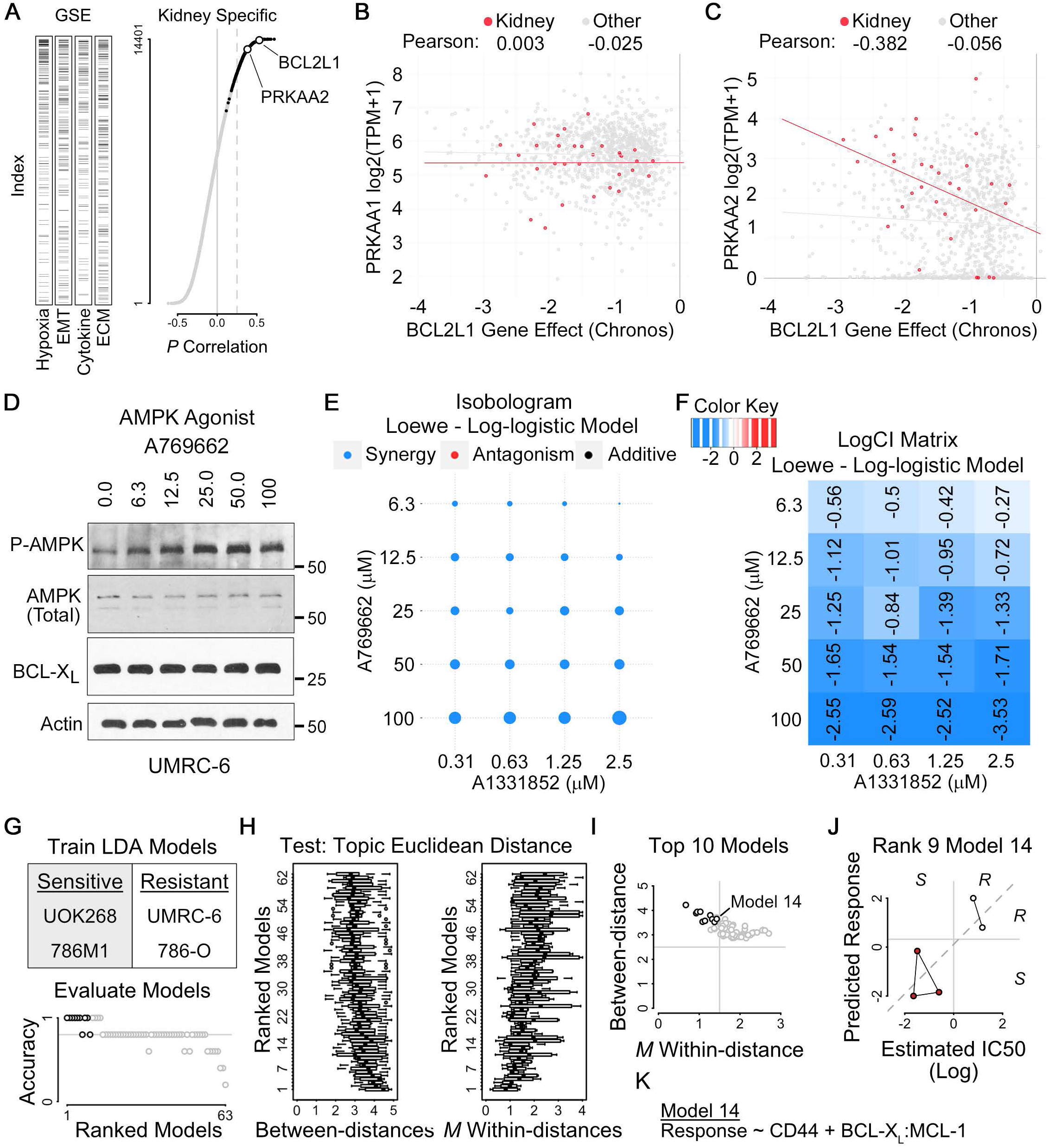
Multivariate Analysis Establish a Biomarker Panel to Predict BCL-X_L_ Dependence. (**A**) Pairwise Pearson correlation values between kidney lineage-specific gene expression and *BCL2L1* dependence (right) and ranked enrichment of the indicated pathways (left; EMT, epithelial-to-mesenchymal transition; ECM, extracellular matrix). Data points in black are correlations which scored as kidney-specific compared to 19 other lineages and 20 quasi-lineages. Correlations were multiplied by -1. (**B** and **C**) Scatter plots comparing gene expression of *PRKAA1* (**B**) and *PRKAA2* (**C**) to *BCL2L1* dependence within the kidney-lineage (red points and line) or all lineages (gray points and line). (**D**) Immunoblot analysis of the indicated proteins in UMRC-6 cells treated with the AMPK agonist, A769662, at the indicated concentrations for 24 hours. (**E** and **F**) Isobologram (**E**) and combination index (CI) plot (**F**) derived from Loewe modeling, using the SiCoDEA algorithm (33), of the percent viability of UMRC-6 cells treated with the indicated concentrations of drug. Blue indicates synergy. (**G**) Linear Discriminate Analysis (LDA) models trained on the indicated cell lines (top) and ranked by their accuracy [true positives (TP) + true negatives (TN) divided by TP + TN + false positives + false negatives] in the test cohort (bottom). Black points are the top 10 ranked models. (**H**) Box-and-Whisker plots of ranked LDA models indicating the between- and mean (*M*) within-Euclidian distances of points as plotted in (**J**). Topics are sensitive or resistant. Euclidian distances were determined by plotting experimentally determined IC50 values versus predicted responses in the test cohort. (**I**) Scatterplot comparing LDA models between-and within-Euclidian distances. Black points are the top 10 ranked models. (**J**) Scatterplot of the estimated IC50 values and the predicted response using model 14. Lines between points connect cell lines in the same topic (sensitive or resistant). Italics on plot margins indicate actual and predicted sensitivity (S) or resistance (R). (**K**) Description of model 14.

*PRKAA2* encodes isoform 2 of the AMP-activated Kinase (AMPK), a well-studied nutrient/energy biosensor (40). Interestingly, many of the features associated with BCL-X_L_ dependence (in studies done by us and others), such as nutrient deprivation, cell cycle exit, metabolic dysfunction, metastatic dormancy, etc., are all associated with activation of AMPK (**Supplementary Fig. S6C**) (41–44). Furthermore, AMPK has been directly linked to changes in BCL-X_L_ dependent apoptosis in liver cancer cells (45). Together, we hypothesized that the seemingly disparate biological programs driving aggressive renal cancer possibly converge of AMPK-dependent control of BCL-X_L_ activity.

Unfortunately, the two isoforms of AMPK are highly conserved and not distinguishable by the best validated antibodies. Consistent with this, we failed to see any notable differences in total AMPK expression levels between the BCL-X_L_i sensitive versus resistant RCC lines (although a smaller migrating band of ∼60 KDa was notable in the BCL-X_L_ dependent A498 and UOK262 cells, **Supplementary Fig. S6D**). Phospho-AMPK was not discernible under standard nutrient-rich growth conditions. Consequently, rather than establishing correlations with AMPK levels, we directly addressed the links between AMPK activation and BCL-X_L_ dependence using functional studies. We treated the BCL-X_L_-inhibitor insensitive UMRC-6 cells with combinations of the AMPK agonist, A769662, and the BCL-X_L_ inhibitor, A1331852. Following 7 days of treatment, we measured the impact of AMPK activation on BCL-X_L_ dependence, using the SiCoDEA algorithm (33). Remarkably, we found that AMPK activation sensitized these cells to BCL-X_L_ loss, showing synergy at various drug combinations (**Figs. 6D** to **6F**). We thus concluded that AMPK activation represents at least one signaling node to sensitize kidney cancer cells to BCL-X_L_ inhibitors.

Finally, we integrated our biomarker studies into a single multivariate panel. Here, we excluded *PRKAA2* (given the caveats described above) and modeled the predictive power of the remaining individual biomarkers. To this end, we chose two representative BCL-X_L_i ‘sensitive’ (i.e., 786M1 and UOK268) versus two ‘resistant’ cells (786O and UMRC-6) and performed ‘linear discriminate analysis (LDA)’ using the protein expression data (**Fig. 6G**). LDA is a method of statistical dimensionality reduction, similar to principal component analysis, which models datasets into separate classes (in this case BCL-X_L_ dependent vs independent). This analysis allowed us to test 63 models, including the univariate and all multivariate *k*-combinations of BCL-X_L_, MCL-1, NOXA, CD44, and ratios of BCL-X_L_:MCL-1 and NOXA:MCL-1. The ideal model was expected to have high accuracy (as measured by predicted IC_50_ value), lower Euclidian distance within the same class, and higher distance between classes (**Figs. 6H** and **6I**). We applied this trained model to the remaining cell lines in our panel. Remarkably, a multivariate model (model 14), representing high CD44 expression and high BCL-X_L_:MCL-1 ratio, reliably predicted the A498, UOK262, and UOK127 cells to be BCL-X_L_i sensitive and the UMRC-2 and UOK276 cells to be BCL-X_L_i insensitive (**Figs. 6J** and **6K**). This clinically adaptable panel offers the opportunity to stratify renal tumors based on BCL-X_L_ dependence.

## Discussion

Our previous studies discovered the relevance of BCL-X_L_ in ccRCC (17); however, the underlying molecular mechanisms of this dependency were unclear. Since then, others have reported the relevance of BCL-X_L_ in kidney cancer cells that have entered dormancy due to therapeutic response (13), or in metastatic settings (18). Altogether, these studies have reinforced the importance of targeting BCL-X_L_ in kidney cancer and highlighted the need to better understand the molecular mechanisms of BCL-X_L_ dependence and the extent to which other (rarer) sub-types of kidney cancer relied on this protein.

In this study, we addressed these conceptual gaps. Exploiting our previous observation that mesenchymal transition promotes BCL-X_L_ dependence, we interrogated the alterations in the BCL-X_L_ interactome and the cellular transcriptome to determine the molecular events associated with higher BCL-X_L_ dependence. These studies revealed a rewiring BCL-X_L_’s protein-protein interaction network in TGFβ treated cells and established that BCL-X_L_ conferred protection against BIM-dependent apoptosis in mesenchymal cells. Additional follow-up studies would, however, be required to fully study the functional implications of this shift in the BCL-X_L_ interactome.

Our studies have consolidated the idea that, besides EMT, cellular features that are typically associated with more aggressive cancers – anoikis resistance and metastasis proficiency – increase cellular dependence on BCL-X_L_. The links between anoikis programs and BCL-X_L_ dependence were revealed by our transcriptomics studies, which also discovered that TGFβ treatment robustly induces the expression of the BCL-family regulatory protein, BMF. Although, BMF loss alone was not sufficient to counteract the effects of TGFβ, it nevertheless revealed how global epithelial ‘detachment’ programs that typically precede metastasis might confer increased BCL-X_L_ dependence. Consistent with this idea we noted that metastatic-proficiency also conferred increased sensitivity to BCL-X_L_ loss. Dormancy programs in mesenchymal/metastatic cells are typically associated with driving chemo- and radiotherapy resistance; however, by activating anoikis-like programs, cancer cells entering dormancy select for BCL-X_L_ function as an anti-apoptotic shield that blocks imminent cell death. This observation is clinically exploitable.

Virtually all of the previous work on the importance of BCL-X_L_ in kidney cancer, including our own studies, have been done in ccRCC. Unfortunately, the rarer kidney cancer subtypes (e.g., translocation RCC, FH-deficient RCC, Renal Medullary Carcinoma, sarcomatoid RCC) are often deadlier than ccRCC. Interestingly, our machine-learning algorithm (TrPLet), which predicts cellular dependencies based on transcriptomics data, revealed that BCL-X_L_, but not the other members of the BCL-2 family, was a subtype agnostic dependency in kidney cancer. We thus addressed the importance of BCL-X_L_ as an oncogenic target in two of the rare kidney cancer models: FH-deficient RCCs and sarcomatoid RCCs. Our studies used physiologically relevant models of these disease types. For example, in our FH studies, we used both traditional isogenic models of this disease but also tested these links using orthogonal approaches where we deployed genetic (CRISPR/Cas9 inactivation), metabolic (MMF administration), and pharmacologic (FH inhibition) approaches to recapitulate the biology of FH loss in pRCC. Likewise, we tested BCL-X_L_ dependence using sarcomatoid models of both ccRCC and chRCC. Remarkably, other than the p53-deficient sarcomatoid chRCC cells, UOK276, which also had very low BCL-X_L_ expression, we noted robust BCL-X_L_ dependence in all other RCC subtypes.

Thrombocytopenia is a major clinical concern associated with the use of BCL-X_L_ inhibitors due to on-target inhibition of BCL-X_L_ in platelets. Interestingly, VHL targeting Proteolysis Targeting Chimeras (PROTACs) do not work as effectively in platelets because of low VHL expression. Importantly, the VHL-engaging BCL-X_L_ PROTAC, DT2216 (46), is already being tested in the clinic (NCT04886622, NCT06620302, and NCT06964009) and could be utilized against FH-deficient RCCs in the future, considering that they retain VHL function. The *VHL*-deficient tumors, instead, can be targeted using cereblon-engaging [e.g., XZ739 (47)], IAP-engaging [e.g., compound **8a** (48)] and/or MDM2-engaging [e.g., BMM4 (49) and AN-1/AN-2 (50)] PROTACs, all of which are also being optimized for platelet-sparing properties, in future clinical studies.

Careful patient stratification strategies, using reliable predictive biomarkers, can also improve therapeutic efficacy. Interestingly, we found that the lineage agnostic biomarkers reported in the DepMap dataset do not faithfully capture the predictors of BCL-X_L_ dependence in kidney cancer. Although, some proteins trended as expected (higher BCL-X_L_, lower MCL1), expression of other proteins (e.g., PMAIP1/NOXA, RBCK1, GTF2IRD1, etc.) showed no correlation with BCL-X_L_ dependence in kidney cancer. One caveat of our analysis, though, was the limited number of cell lines available for these studies, especially in the non-ccRCC subtypes. Nevertheless, applying both univariate and multivariate models, we interrogated lineage-specific biomarkers for RCC and developed a predictive panel that could accurately demarcate cells based on BCL-X_L_ dependence, agnostic to the RCC subtype. These studies revealed the importance of high *PRKAA2*, which encodes AMPKα2 as a driver of BCL-X_L_ dependence in kidney cancer. Our findings suggest that seemingly disparate biological programs (e.g., nutrient limitation, metastatic dormancy, therapeutic senescence, etc.) could converge on AMPK to control BCL-X_L_-dependent cell death programs. However, although AMPKα1 and AMPKα2 are highly conserved proteins, only AMPKα2 expression correlates with BCL-X_L_ dependence, suggesting a context-dependent division of labor between these proteins. These links will be actively pursued in future studies.

Undesirable off-target effects represent one of the major concerns of pharmacological studies. In this regard, we relied on the use of well-annotated antagonists of the BCL-2 family proteins, including A1331852, which is a highly selective BCL-X_L_ inhibitor (51, 52).

Additionally, in our previous work, we undertook extensive studies to establish that the cytotoxic effects of A1331852 were on-target, including the use of drug-resistant mutants to reverse these fitness defects in kidney cancer (17). In these earlier studies, we also performed in vivo studies in mouse ccRCC models to demonstrate the efficacy of BCL-X_L_ blockade. Here, we extended these findings by using a machine learning model in human tumor datasets, validated these predictions in physiologically relevant cell lines, and ultimately addressed the relevance of BCL-X_L_ blockade in metastatic disease.

Our studies, alongside a growing body of work reported by others, identifies the oncogenic relevance of BCL-X_L_ in kidney cancer. Our work has clinical relevance across a range of RCC subtypes, including some of the rare and aggressive forms of this disease. The increased mechanistic understanding and biomarker discovery in this work offers an actionable path to transition our findings into the clinic and justifies the need to target BCL-X_L_, especially in aggressive/metastatic forms of kidney cancer. With already growing attention to platelet-sparing BCL-X_L_ degraders in the clinic, our work establishes the opportunity to rapidly repurpose these agents into RCC therapy.

## Acknowledgments/COIs

The authors declare no conflicts of interest associated with this study. This work was primarily supported by the DOD-KCRP-IDA award (HT9425-23-1-0771) to AAC and KAS. AAC was additionally supported by seed money by the Cleveland Clinic Foundation, pilot grants by the Case Comprehensive Cancer Center, DOD-KCRP-ECI award (W81XWH-20-1-0804), DOD-KCRP-IDA (HT9425-24-1-0651), DOD-KCRP-IDA (HT9425-25-1-0761), two Velosano pilot awards, the V foundation scholar award (V2020-011), the ACS-Research Scholar Grant (RSG-22-067-01-TBE), and the NCCN-YIA. KAS was supported by NIH R37CA248565, NIH R01DK125263, Alex’s Lemonade Stand Foundation, Andrew McDonough B+ Foundation, Blavatnik Institute at Harvard, and Ovarian Cancer Research Alliance grant CRDG-2023-3-1001.

## Materials and Methods

### Cell Lines

A-498 (RRID: CVCL_1056), 786-O (RRID: CVCL1051), and ACHN (RRID:CVCL_1067) renal carcinoma cells were obtained from American Type Culture collection. HEK293T (RRID: CVCL_0063) was obtained from the Lerner Research Institute’s Cell Media Core. Above cells were maintained in Dulbecco’s Modified Eagles Medium (DMEM) (Life Technologies 11995073) supplemented with 10% Fetal Bovine Serum (FBS) (Life Technologies 10437-028), and 1X Penicillin-Streptomycin (Life Technologies 15140163). UOK262 (UOK262^FH-null^) and UOK268 (UOK268^FH-null^) and their isogenic FH-proficient counterparts (UOK262^FH-WT^ and UOK268^FH-WT^) created by the stable reintroduction of the wild-type *FH* gene back into the established cell lines, and the sarcomatoid RCCs, UOK127 and UOK276, were all obtained as a generous gift from Dr. Marston Linehan (National Cancer Institute). These cells were maintained in Dulbecco’s Modified Eagles Medium (DMEM) (Life Technologies 11995073) supplemented with 10% Heat inactivated Fetal Bovine Serum (HI-FBS) (Gibco 5256801), 1X Penicillin-Streptomycin (Life Technologies 15140163), non-essential amino acid (NEAA) supplement (Gibco 11140050). Cells were incubated at 37°C in a humidified environment with 5% CO₂. Where relevant, cells were selected using Puromycin (2 µg/ml) for 3-5 days or Blasticidin (10 µg/ml) for 7-10 days.

### Plasmids

CRISPR/Cas9 constructs were generated using restriction enzyme-based cloning. BsmBI-digested pLentiCRISPRv2 plasmid (RRID: Addgene_52961) was ligated to annealed oligos encoding sgRNAs targeting *BMF* or *FH*. sgRNA sequences were designed using CRISPick (RRID: SCR_025148). All plasmids were confirmed by Sanger Sequencing.

### Lentiviral Infections

Lentivirus was generated using HEK293T cells. 1.8E06 to 2.6E06 293T cells were seeded into 6 cm plates. The following day, cells were transfected using 3 μg of DNA [1.5 µg lentiviral plasmid + 1.5 µg helper plasmids - psPAX2 (RRID: Addgene_12260) and pMD2.G (RRID: Addgene_12259) mixed in a 3:1 ratio], combined with 9 µl lipofectamine 2000 (Invitrogen 11668019) in Opti-MEM Reduced Serum media (Gibco, 31985062). After incubation at room temperature for 15 minutes, transfection mixtures were added dropwise onto cells and incubated for additional 6-8 hours at 37 °C in a humidified environment with 5% CO₂. Media was changed, and viral supernatant was collected at 48- and 72-hours post transfection. Lentiviral supernatants were combined and filtered (0.45 μm). 1.5E05 to 2.0E05 cells were seeded per well into 6-well plates and transduced with 0.5 mL of lentiviral supernatant in presence 2 μg/mL polybrene. Plates were centrifuged at 300× *gg* for 40 min and returned to the incubator. After 8 hours media was replaced. Transduced cells were grown for an additional 24 hours prior to adding selection (2 µg/ml puromycin).

### Flow cytometry based BH3 Profiling

BH3 profiling was performed by flow cytometry as previously described (23). Briefly, cells were collected and resuspended in Mannitol Experimental Buffer (MEB), consisting of 10 mM HEPES (pH 7.5), 150 mM mannitol, 50 mM KCl, 0.02 mM EGTA, 0.02 mM EDTA, 0.1% BSA, and 5 mM succinate. The resuspended cells were subsequently added into pre-prepared assay plates containing the indicated BH3 peptides and 0.001% digitonin. Alamethicin (25 μM) was used as positive control, while DMSO was used as negative control. After incubation at 28°C for 60 minutes, cells were fixed with paraformaldehyde for 15 minutes, followed by neutralization with N2 buffer (1.7 M Tris base, 1.25 M glycine, pH 9.1). Fixed cells were then stained overnight at 4 °C in intracellular staining buffer (0.2% Tween-20, 1% BSA) containing DAPI (1:1000, Abcam) and anti-Cytochrome c-Alexa Fluor 647 (1:2000, clone 6H2.B4, BioLegend). Cytochrome c release was quantified using an Attune NxT flow cytometer (Thermo Fisher Scientific).

### RNA isolation and quantitative real-time PCR

Total RNA was purified using TRIzol^TM^ (ThermoFisher, 15596026), as per the manufacturer’s instructions, and reverse transcribed into cDNA using the Reverse Transcription kit (Qiagen). Quantitative PCR (qPCR) was performed in SYBR Green master mix (Qiagen) with an Applied Biosystems 7500 PCR. The mRNA levels were normalized to *β-Actin*.

### Real-time PCR Oligonucleotide Primer sequences

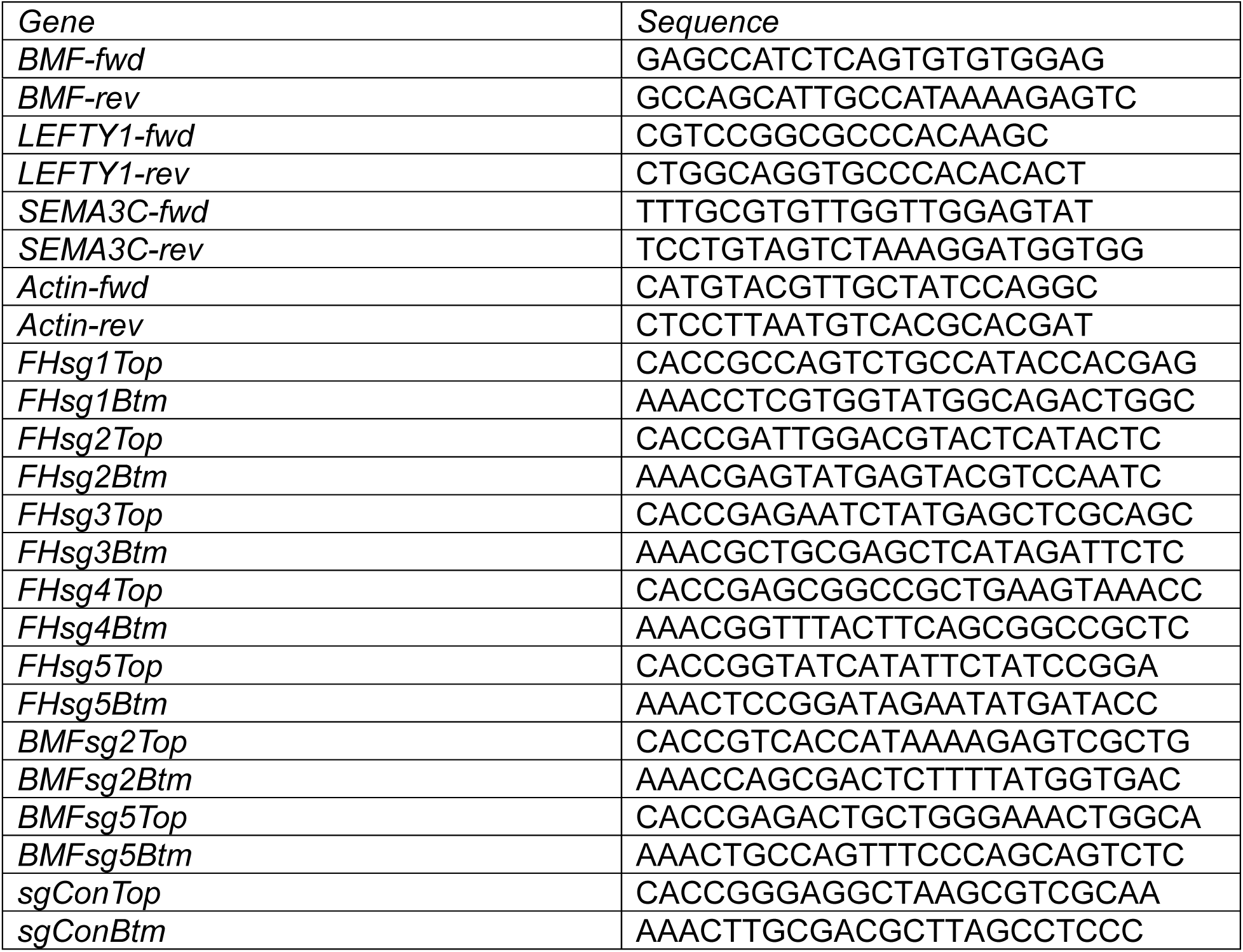

### Immunoprecipitation and LC-MS. Immunoprecipitation and LC-MS

UMRC-2 cells (∼2-3 x 15 cm plates; ∼3.0E07 cells) treated with either DMSO or TGFβ were harvested and washed once in 1X PBS. Pellets were resuspended in ice-cold lysis buffer, centrifuged at 8000xg for 10 minutes at 4°C. Including a negative control (Normal Rabbit IgG), pre-clearing of lysates were done using Agarose A/G beads to precipitate unwanted proteins and increase purity of proteins and reduce non-specific binding.

Samples derived from IgG and BCL-X_L_ pulldown experiments were digested with trypsin to be analyzed by LC-MS/MS. Immunoprecipitated material was subjected to on-bead tryptic digestion (53). Briefly, trypsin (10 µl, 10 ng/µl) in 100 mM ammonium bicarbonate was added to washed beads, and samples vortexed at room temperature for 15 sec every 2–3 min for 15 min to ensure even suspension of beads. Digestion was carried out overnight at 37°C. A second 10-µl aliquot of protease was added for 4 h at 37°C. After digestion, the samples were centrifuged, and the supernatant was collected and diluted with formic acid (5% v/v final concentration). The digests were cleaned using PepClean C-18 spin columns (ThermoFisher) according to manufacturer’s instructions, the samples dried in a vacuum concentrator and reconstituted in 0.1% formic acid.

The Mass spectrometry analysis of the tryptic digests was performed on a Orbitrap Exploris 480 mass spectrometer equipped with a Vanquish Neo uHPLC system (Thermo Fisher Scientific, Waltham, MA). The HPLC column was a 50 cm x 75 µm id Easy Spray Pepmap NEO C18, 2μm, 100 Å reversed-phase capillary chromatography column. 5 μL volumes of the peptide samples were injected, eluted from the column by an ACN/0.1% FA gradient at a flow rate of 0.3 μL/min and were introduced into the source of the mass spectrometer on-line. Digests were analyzed using the data-dependent multitask capability of the instrument, acquiring full-scan mass spectra from 350-1400 Da (120k resolution) to determine peptide molecular weights and product ion spectra acquired using HCD (30%nce, 15k resolution) to determine amino acid sequence in successive instrument scans. The Exploris data was analyzed by using all HCD spectra collected in the experiment to search the human SwissProt database (Downloaded on 3-23-2022, 20576 entries) using Proteome Discoverer 2.5. The parameters used for these searches include methionine oxidation and phosphorylation (S,T,Y) as variable modifications, protein mass tolerance of 10 ppm, and fragment ion mass tolerance of 0.06 Da. Protein and peptide validation was performed using Percolator with application of a 1% false discovery rate (FDR) to identify high-confidence proteins.

The relative abundance of the proteins identified in the IgG and BCL-X_L_ was determined using Label free quantitation which was performed using the Minora node in PD2.5. Quantitation was performed on unique peptides and only proteins identified by 2 peptides were quantified and the LFQ intensities were normalized to total peptide content for each LC-MS experiment.

Data was searched against human SwissProtKB database. Over 490 proteins were identified in these samples with the most abundant components corresponding to Actin, Vimentin, Stress-70 protein, Endoplasmic reticulum chaperone BiP, Heterogeneous nuclear ribonucleoproteins A2/B1, Myosin-9, Spectrin, and Tropomyosin alpha-4 chain. The bait protein, BCL-X_L_, was identified as a major component in the BCL-X_L_ IP samples. A comparison of the relative abundance of the recovered proteins from the DMSO and TGFβ samples was used for further studies.

### RNA-Seq Analysis

Total RNA was extracted from the 786-0, 786M1, UMRC-2 (DMSO) and UMRC-2 (TGFβ treated) ccRCC cell lines using TRIzol^TM^ (Life Technologies), quantified using the Qubit RNA Assay Kit (Life Tech), and quality was determined on the Bioanalyzer using the RNA Pico Kit (Agilent). cDNA libraries were prepared and prepared DNA libraries which passed all QC criteria were sequenced as previously described [1]. Raw sequences (.fastq files) were QC tested and total read counts per gene were measured and normalized. Differential expressions were determined using EdgeR (v3.12). GSEA was performed on the EdgeR normalized counts, using downloaded software (http://www.broad.mit.edu/gsea/downloads.jsp), where gene sets with normalized enrichment score >1.5, FDR<0.10 and a nominal *p*<0.05 were considered significant.

### Protein extraction and immunoblot analysis

Cells were rinsed with ice-cold 1XPBS, scraped, and lysed on ice using 1X RIPA buffer (BIORAD) or home-made lysis buffer [50 mM Tris.Cl (pH 7.5), 400 mM NaCl, 1% Nonidet P-40, 1 mM EDTA, and 10% glycerol] supplemented with Protease Inhibitor Cocktail tablet for 30 mins at 4_°_C. Extracted proteins were quantified using the Bradford Assay (Biorad 5000006) and ∼30-50 μg protein was analyzed by SDS-PAGE. For blots with multiple ccRCC cell lines, membranes were stained with Ponceau S solution (Sigma P3504), RT/ 5 mins and imaged before immunoblotting. The following primary antibodies (from Cell Signaling Technology, unless otherwise noted) were used at a 1:1000 dilution: β-Actin (13E5) Rabbit mAb (4970), CD44 (E7K2Y0 XP Rabbit mAb (37259), BCL-X_L_ Antibody (2762), MCL-1 (D2W9E) Rabbit mAb (PE Conjugate, 65617), Vinculin (E1E9V) XP^®^ Rabbit mAb (13901), Fumarase (D9C5) Rabbit mAb (4567), HIF2α (E8E5Z) Rabbit mAb (71565), BMF (E5U2J) Rabbit mAb (50542), Noxa (D8L7U) Rabbit mAb (14766), GTF2IRD1 Ab (Abcam 64805), RBCK1 antibody (Abcam, EPR28157-53), and CD24 (Proteintech 10600-1-AP). Primary antibodies were detected with HRP-conjugated secondary antibodies (Pierce, 1:2000) and chemiluminescent HRP substrates including either Luminol Pierce 32106, Pierce ECL Plus Thermo Fisher Scientific or Millipore Immobilon ECL Ultra.

### Cell Viability Assays

ccRCC lines were seeded overnight (UMRC-2, 786-0, 786M1, TK10, UOK262^FH-null^, UOK268^FH-null^, UOK262^FH-WT^ and UOK268^FH-WT^: 100 cells/well; ACHN: 500 cells/well; in 96-well plates) and then treated with the respective inhibitors or vehicle DMSO. Cells were exposed for 7 days to a two-fold serial dilution of A-1331852 and ABT-199 (Venetoclax) from 0 to 5 μM. Cell viability was analyzed using either XTT assay (Cell Proliferation Kit II, Roche) or CellTiter-Glo (Promega) following manufacturer’s instructions. IC_50_ values were determined using Graphpad Prism (RRID:SCR_000306). In experiments comparing the drug response curves between two (or more) conditions, we used the worksheet provided alongside a previously described statistical method that uses a modified Chi-square analysis to measure significance, regardless of the functionality of the curve (22). Drug-drug interactions were studied by treating cells with a matrix of drug combinations, following which cell viability was measured using the XTT assay. The resulting data was processed and analyzed using the ‘Single and Combined Drug Effect Analysis (SiCoDEA)’ algorithm (33), following the authors’ instructions.

### In Vivo Studies

Rodent (NSG male mice) experiments were performed following all Cleveland Clinic IACUC (protocol no. 0002168) guidelines. For the metastatic assay, 300K luciferin expressing 786M1 cells were injected in a 100 µl volume into the lateral caudal vein of NSG mice. Initial lung engraftment was evaluated 2-days post injection. Following ∼20 days, mice were placed on vehicle or A-1331852 treatment BID. A-1331852 was purchased from MedChemExpress (Catalog: HY-19741). Formulations of A-1331852 (25 mg/kg) were prepared in 60% (v/v) Phosal 50PG, 27.5% (v/v) PEG400, 10% (v/v) ethanol and 2.5% (v/v) DMSO, and oral gavage was done twice a day for up to 4 weeks. ROI were measured weekly using IVIS. Kidney weights or lung tumors were quantified at termination of the experiment. Tumor and normal tissues were harvested upon IHC and CC3 analyses.

### Flow cytometry based BH3 Profiling

The ccRCC cells, pre-treated with TGFβ or Vehicle control for 4 days, were plated in 96-well plates and permeabilized using Mannitol Experimental Buffer (MEB) containing digitonin. Peptides were prepared at appropriate concentrations and loaded into wells, followed by addition of cells. After incubation, cells were stained with antibodies for cytochrome C and DAPI and incubated overnight at 4°C. Flow cytometry was used to measure cytochrome C release for quantitation of mitochondrial outer membrane permeabilization. Controls, including Alamethicin, recombinant caspase-3, and apoptotic inhibitors were included as controls to measure apoptotic priming and dependencies.

### Modeling BCL-X_L_ Dependence

BCL-X_L_ dependence was modeled in R (version 4.5.0) using Linear Discriminate Analysis (MASS, version 7.3-65). The dependent variable [sensitive (S) or resistant (R)] was determined using estimated IC_50_ values for each cell line. A-498, 786M1, UOK127, UOK262, and UOK268 (which represent cell lines with low nanomolar range A1331852 IC50 values) were called sensitive and UOK276, 786-O, UMRC-2, and UMRC-6 (which represent cell lines with high micromolar A1331852 IC_50_ values or undetermined IC_50_ values) were called resistant. Independent variables were calculated using the relative protein expression of BCL-X_L_, MCL-1, NOXA, and CD44. Models were trained on 2 sensitive lines (UOK268 and 786M1) and 2 resistant lines (786-O and UMRC-6). Random assignment to training or testing cohorts was performed a single time at the beginning of the analysis. We trained 63 models using univariate and all multivariate *k*-combinations of independent predictor variables (relative levels of BCL-X_L_, MCL-1, NOXA, CD44, and ratios of BCL-X_L_:MCL-1 and NOXA:MCL-1). Trained models were tested on the remaining cell lines [3 sensitive cell lines (A-498, UOK127, and UOK262) and 2 resistant cell lines (UOK276 and UMRC-2)]. To compare model performance, we calculated model accuracy and used Euclidean distances to determine topic withinness and betweenness. Euclidean distances were determined by plotting the predicted responses and log-transformed IC50 values for each model. We prioritized models with 1) high accuracy, 2) low mean topic withinness, 3) high topic betweenness, and 4) relatively small model coefficients. Models were ranked using the following method.

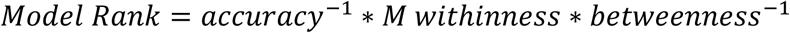

### Determining correlations between kidney lineage-specific gene expression and BCL2L1 dependence

Cell line transcripts per million for all human protein coding genes and CRISPR gene effect scores were obtained from the DepMap. There were a total of 1089 cell lines with both gene expression and *BCL2L1* dependence data. For each of the 19,215 transcriptionally profiled genes, we determined lineage-agnostic and -specific pairwise Pearson correlation values between gene expression and *BCL2L1* dependence. For lineage-agnostic correlations, we used all 1089 cell lines. For lineage-specific correlations, we excluded lineages which had less than 20 cell lines to ensure sufficient representation. This threshold left 20 lineages for comparison. Genes which showed lineage-specificity were determined by taking the lineage-specific Pearson correlation values for each gene and converting them into Z-scores. To improve robustness, Z-scores were calculated using each lineage and 20 additional randomly generated quasi-lineages with a matching distribution of sample sizes. Genes which 1) had a Z-score ≤-2 in the kidney lineage, 2) did not have a Z-score ≤-2 in the bowel lineage (the bowel lineage also displays *BCL2L1* dependence and in this analysis was used as a control), and 3) had an average kidney lineage-specific transcript per million >0.25 were called kidney-specific. Gene set enrichment was performed on the ranked kidney-specific Pearson correlation values by calculating a running-sum statistic (enrichment) for each gene set. Significance was estimated by comparing the maximum enrichment to 100 randomly generated gene set permutations of matching size.

## References

1. Siegel, R.L., et al., Cancer statistics, 2025. CA Cancer J Clin, 2025. 75(1): p. 10–45.

2. Srinivasan, R., et al., New strategies in renal cell carcinoma: targeting the genetic and metabolic basis of disease. Clin Cancer Res, 2015. 21(1): p. 10–7.

3. Linehan, W.M., et al., Genetic basis of cancer of the kidney: disease-specific approaches to therapy. Clin Cancer Res, 2004. 10(18 Pt 2): p. 6282S–9S.

4. Ricketts, C.J., et al., The Cancer Genome Atlas Comprehensive Molecular Characterization of Renal Cell Carcinoma. Cell Rep, 2018. 23(1): p. 313–326 e5.

5. Schmidt, L., et al., Germline and somatic mutations in the tyrosine kinase domain of the MET proto-oncogene in papillary renal carcinomas. Nat Genet, 1997. 16(1): p. 68–73.

6. Delahunt, B., Sarcomatoid renal carcinoma: the final common dedifferentiation pathway of renal epithelial malignancies. Pathology, 1999. 31(3): p. 185–90.

7. Prado, L.C., et al., Management of Renal Cell Carcinoma of Variant Histology. JCO Oncol Pract, 2025: p. OP2500640.

8. Yu, Y., et al., Comparative analysis of clinicopathological characteristics and prognostic outcomes in fumarate hydratase-deficient renal cell carcinoma versus high-grade papillary renal cell carcinoma. Transl Androl Urol, 2025. 14(8): p. 2195–2206.

9. Wang, Y.S., et al., Analysis of the factors influencing the survival time of patients with sarcomatoid renal cell carcinoma. Mol Clin Oncol, 2019. 11(4): p. 405–410.

10. Boise, L.H., et al., bcl-x, a bcl-2-related gene that functions as a dominant regulator of apoptotic cell death. Cell, 1993. 74(4): p. 597–608.

11. Singh, R., A. Letai, and K. Sarosiek, Regulation of apoptosis in health and disease: the balancing act of BCL-2 family proteins. Nat Rev Mol Cell Biol, 2019. 20(3): p. 175–193.

12. Zhou, L., et al., P21-positive senescent stromal cells promote prostate cancer immune suppression and progression that can be reversed by senolytic therapy. Cancer Discov, 2025.

13. Xue, Y., et al., Osalmid sensitizes clear cell renal cell carcinoma to navitoclax through a STAT3/BCL-XL pathway. Cancer Lett, 2025. 613: p. 217514.

14. Alcon, C., et al., HRK downregulation and augmented BCL-xL binding to BAK confer apoptotic protection to therapy-induced senescent melanoma cells. Cell Death Differ, 2025. 32(4): p. 646–656.

15. Sela, Y., et al., Bcl-xL Enforces a Slow-Cycling State Necessary for Survival in the Nutrient-Deprived Microenvironment of Pancreatic Cancer. Cancer Res, 2022. 82(10): p. 1890–1908.

16. Soderquist, R.S., et al., Systematic mapping of BCL-2 gene dependencies in cancer reveals molecular determinants of BH3 mimetic sensitivity. Nat Commun, 2018. 9(1): p. 3513.

17. Grubb, T., et al., A Mesenchymal Tumor Cell State Confers Increased Dependency on the BCL-XL Anti-apoptotic Protein in Kidney Cancer. Clin Cancer Res, 2022.

18. Schoenfeld, D.A., et al., Immune dysfunction revealed by digital spatial profiling of immuno-oncology markers in progressive stages of renal cell carcinoma and in brain metastases. J Immunother Cancer, 2023. 11(8).

19. Tsherniak, A., et al., Defining a Cancer Dependency Map. Cell, 2017. 170(3): p. 564–576 e16.

20. Mahmoud, N., et al., BCL-xL dependency in chromophobe renal cell carcinoma. Cancer Gene Ther, 2025. 32(10): p. 1133–1143.

21. Sadagopan, A., et al., Cancer target discovery enabled by transcriptome-based virtual CRISPR screening. bioRxiv, 2025.

22. Hristova, K. and W.C. Wimley, Determining the statistical significance of the difference between arbitrary curves: A spreadsheet method. PLoS One, 2023. 18(10): p. e0289619.

23. Fraser, C., J. Ryan, and K. Sarosiek, BH3 Profiling: A Functional Assay to Measure Apoptotic Priming and Dependencies. Methods Mol Biol, 2019. 1877: p. 61–76.

24. Huber, M.A., N. Kraut, and H. Beug, Molecular requirements for epithelial-mesenchymal transition during tumor progression. Curr Opin Cell Biol, 2005. 17(5): p. 548–58.

25. Subramanian, A., et al., Gene set enrichment analysis: a knowledge-based approach for interpreting genome-wide expression profiles. Proc Natl Acad Sci U S A, 2005. 102(43): p. 15545–50.

26. Liberzon, A., A description of the Molecular Signatures Database (MSigDB) Web site. Methods Mol Biol, 2014. 1150: p. 153–60.

27. Puthalakath, H., et al., Bmf: a proapoptotic BH3-only protein regulated by interaction with the myosin V actin motor complex, activated by anoikis. Science, 2001. 293(5536): p. 1829–32.

28. Vanharanta, S., et al., Epigenetic expansion of VHL-HIF signal output drives multiorgan metastasis in renal cancer. Nat Med, 2013. 19(1): p. 50–6.

29. Eng, C., et al., A role for mitochondrial enzymes in inherited neoplasia and beyond. Nat Rev Cancer, 2003. 3(3): p. 193–202.

30. Sciacovelli, M., et al., Fumarate is an epigenetic modifier that elicits epithelial-to-mesenchymal transition. Nature, 2016. 537(7621): p. 544–547.

31. Bear, A., et al., Characterization of two human cell lines (TK-10, TK-164) of renal cell cancer. Cancer Res, 1987. 47(14): p. 3856–62.

32. Takeuchi, T., P.T. Schumacker, and S.A. Kozmin, Identification of fumarate hydratase inhibitors with nutrient-dependent cytotoxicity. J Am Chem Soc, 2015. 137(2): p. 564–7.

33. Spinozzi, G., et al., SiCoDEA: A Simple, Fast and Complete App for Analyzing the Effect of Individual Drugs and Their Combinations. Biomolecules, 2022. 12(7).

34. Yang, Y., et al., UOK 262 cell line, fumarate hydratase deficient (FH-/FH-) hereditary leiomyomatosis renal cell carcinoma: in vitro and in vivo model of an aberrant energy metabolic pathway in human cancer. Cancer Genet Cytogenet, 2010. 196(1): p. 45–55.

35. Yang, Y., et al., A novel fumarate hydratase-deficient HLRCC kidney cancer cell line, UOK268: a model of the Warburg effect in cancer. Cancer Genet, 2012. 205(7-8): p. 377–90.

36. He, H. and C. Magi-Galluzzi, Epithelial-to-mesenchymal transition in renal neoplasms. Adv Anat Pathol, 2014. 21(3): p. 174–80.

37. Gnarra, J.R., et al., Mutations of the VHL tumour suppressor gene in renal carcinoma. Nature Genetics, 1994. 7(1): p. 85–90.

38. Yang, Y., et al., Genomic and metabolic characterization of a chromophobe renal cell carcinoma cell line model (UOK276). Genes Chromosomes Cancer, 2017. 56(10): p. 719–729.

39. Grubb, T., et al., The SLC1A1/EAAT3 dicarboxylic amino acid transporter is an epigenetically dysregulated nutrient carrier that sustains oncogenic metabolic programs. Nat Commun, 2025. 16(1): p. 10012.

40. Woods, A., et al., The alpha1 and alpha2 isoforms of the AMP-activated protein kinase have similar activities in rat liver but exhibit differences in substrate specificity in vitro. FEBS Lett, 1996. 397(2-3): p. 347–51.

41. Kemp, B.E., et al., Dealing with energy demand: the AMP-activated protein kinase. Trends Biochem Sci, 1999. 24(1): p. 22–5.

42. Narbonne, P. and R. Roy, Inhibition of germline proliferation during C. elegans dauer development requires PTEN, LKB1 and AMPK signalling. Development, 2006. 133(4): p. 611–9.

43. Karpel-Massler, G., et al., Induction of synthetic lethality in IDH1-mutated gliomas through inhibition of Bcl-xL. Nat Commun, 2017. 8(1): p. 1067.

44. Penugurti, V., et al., AMPK: The energy sensor at the crossroads of aging and cancer. Semin Cancer Biol, 2024. 106**-**107: p. 15–27.

45. Yang, D., et al., Adenosine activates AMPK to phosphorylate Bcl-XL responsible for mitochondrial damage and DIABLO release in HuH-7 cells. Cell Physiol Biochem, 2011. 27(1): p. 71–8.

46. Khan, S., et al., A selective BCL-X(L) PROTAC degrader achieves safe and potent antitumor activity. Nat Med, 2019. 25(12): p. 1938–1947.

47. Zhang, X., et al., Discovery of PROTAC BCL-XL degraders as potent anticancer agents with low on-target platelet toxicity. Eur J Med Chem, 2020. 192: p. 112186.

48. Zhang, X., et al., Discovery of IAP-recruiting BCL-XL PROTACs as potent degraders across multiple cancer cell lines. Eur J Med Chem, 2020. 199: p. 112397.

49. Chang, M., et al., MDM2-BCL-X(L) PROTACs enable degradation of BCL-X(L) and stabilization of p53. Acta Mater Med, 2022. 1(3): p. 333–342.

50. Liu, M., et al., Synthesis of selective BCL-X L PROTAC and potent antitumor activity in glioblastoma. Cell Reports Physical Science, 2025. 6(5).

51. Wang, L., et al., Discovery of A-1331852, a First-in-Class, Potent, and Orally-Bioavailable BCL-XL Inhibitor. ACS Med Chem Lett, 2020. 11(10): p. 1829–1836.

52. Leverson, J.D., et al., Exploiting selective BCL-2 family inhibitors to dissect cell survival dependencies and define improved strategies for cancer therapy. Sci Transl Med, 2015. 7(279): p. 279ra40.

53. Mohammed, H., et al., Rapid immunoprecipitation mass spectrometry of endogenous proteins (RIME) for analysis of chromatin complexes. Nat Protoc, 2016. 11(2): p. 316–26.

