## Supplementary Figures for "BCL-X_L_ Dependence is a Subtype Agnostic Actionable Feature of Difficult-to-Treat Kidney Cancers"

*Supplementary Figure S1*

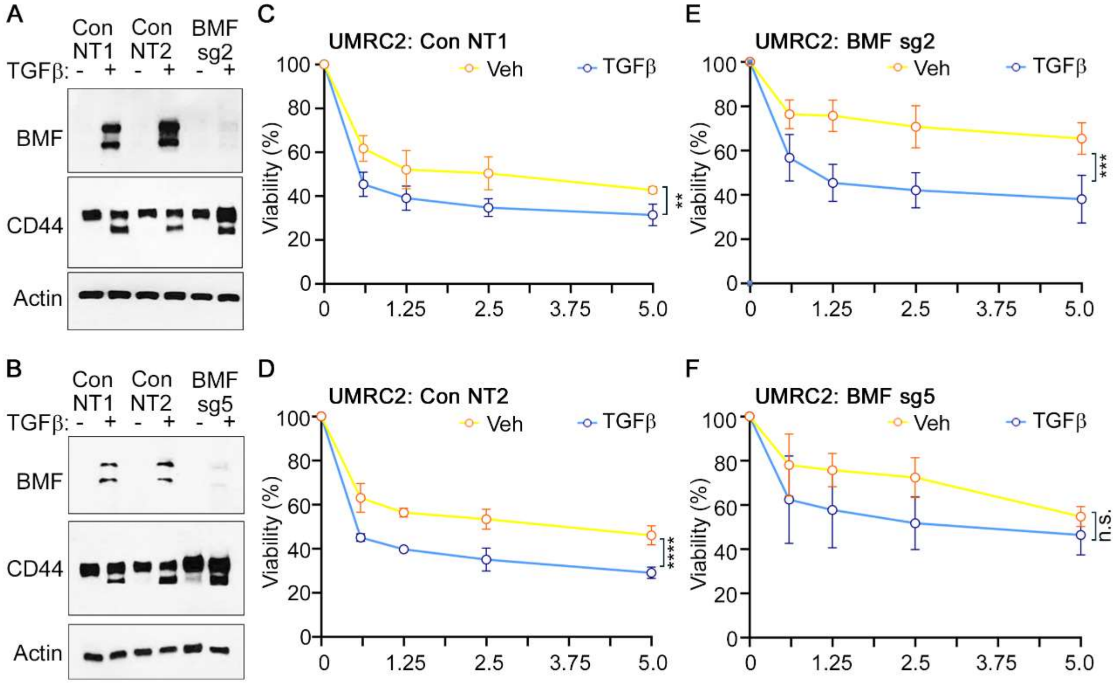


**Supplementary Figure S1. BMF Inactivation is Not Sufficient to Counteract TGFβ Dependent Sensitization to BCL-XL Blockade.** (**A** and **B**) Immunoblot analysis of the indicated proteins in UMRC-2 cells lentivirally transduced to express sgRNAs targeting *BMF* (sg2 and sg5) or non-targeting sgRNAs (NT1 and NT2) and treated with or without TGFβ (20 μg/ml) for 4-days. (**C** to **F**) Percent viability of cells as described in (**A**) and (**B**) treated with increasing concentrations (μM) of A1331852 for 5-7 days. Curves were compared using the modified Chi-square, curve fit analysis (21); n=3, **p<0.01, ***p<0.001, ****p<0.0001, n.s.=not-significant.

*Supplementary Figure S2*

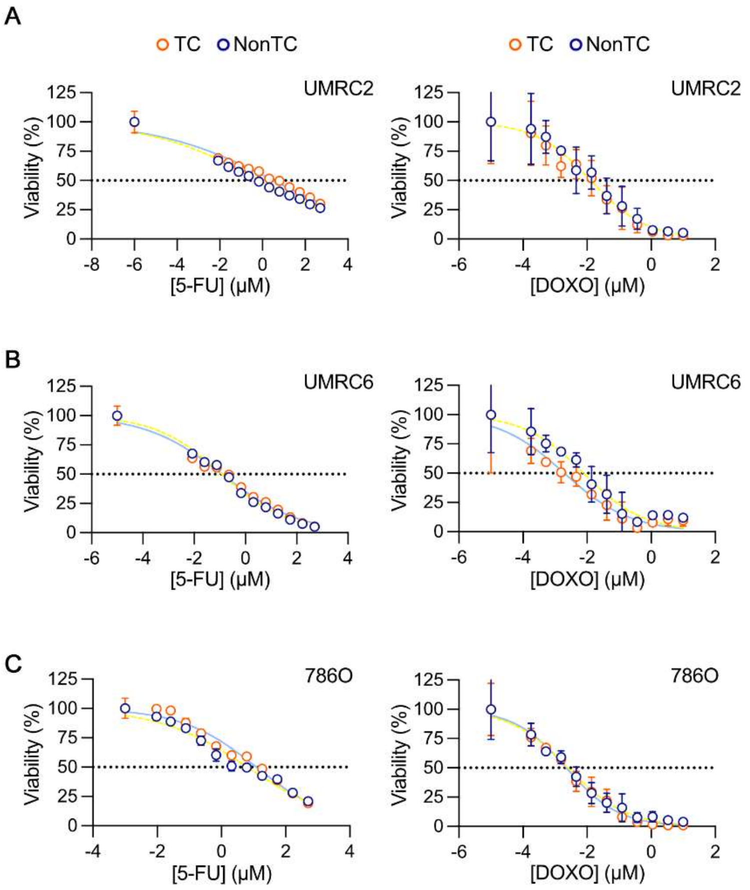


**Supplementary Figure S2. Detached Culture Conditions Do Not Sensitize ccRCC Cells to Standard Chemotherapeutics.** (**A** to **C**) Percent cell viability of the UMRC-2 (**A**), UMRC-6 (**B**), and 786O (**C**) cell lines, measures using CellTiter-Glo, after treatment with serial dilutions of either 5-flurouracil (5-FU) or Doxorubicin (DOXO) for 3-days. Drug concentrations are log-transformed.

*Supplementary Figure S3*

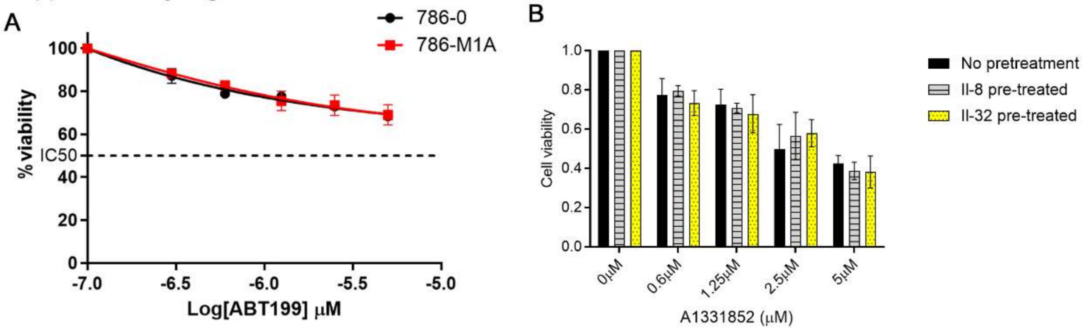


**Supplementary Figure S3. Cytokine Administration is Not Sufficient to Induce BCL-$X_L$ in Non-metastatic cells.** (**A**) Percent viability of the indicated cell lines, as measured using CellTiter-Glo, after treatment with the indicated concentrations of ABT199 for 5-7 days. (**B**) Percent viability of 786-O cells, as measured using CellTiter-Glo, upon pretreatment with the indicated cytokines and then cultured in the presence of the indicated concentrations of A1331852 for 5-7 days.

Supplementary Figure S4

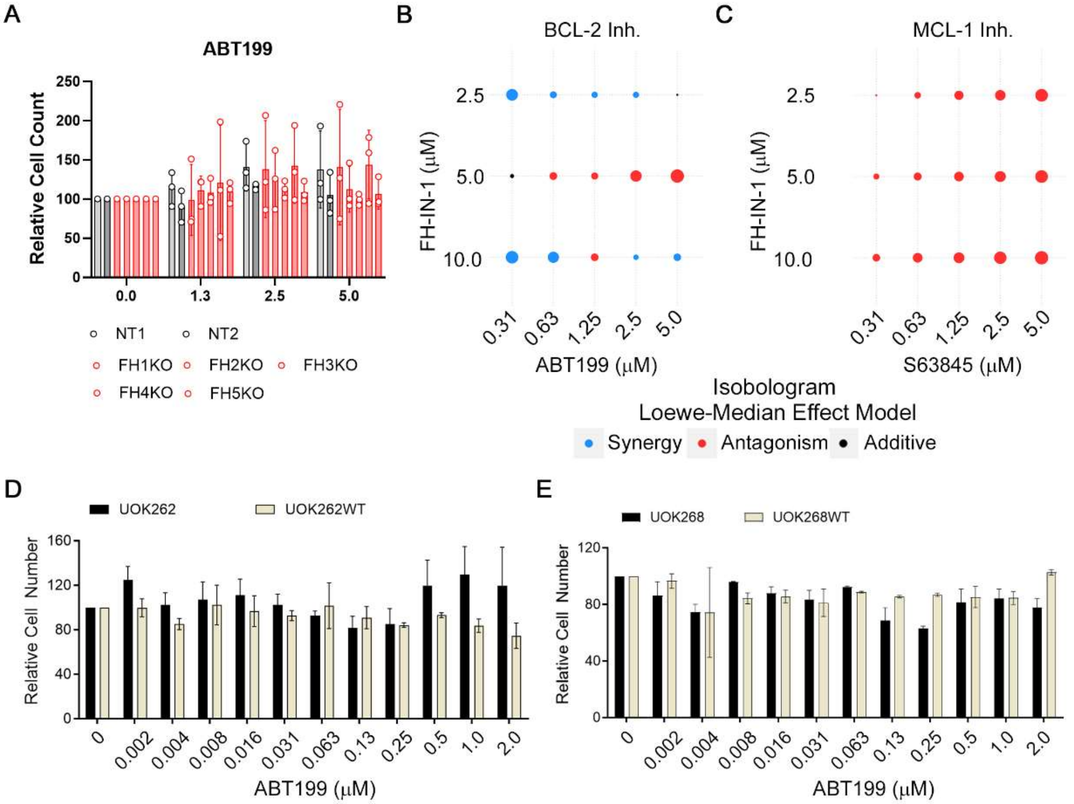


**Supplementary Figure S4. Fumarate Hydratase Inactivation Does Not Impact Response to BCL-2 and MCL-1 Blockade.** (**A**) Relative cell counts of TK-10 cells, as measured using CellTiter-Glo, in cells that were lentivirally transduced to express sgRNAs targeting fumurate hydratase (FH) or non-targeting controls (NT), as indicated, and then treated with the indicated concentrations (µM) of ABT199. (**B** and **C**) Isobolograms derived from Loewe modeling, using the SiCoDEA algorithm (32), of the percent viability of ACHN cells that were treated with the indicated drug combinations of the FH inhibitor (FH-IN-1) with either the BCL-2 inhibitor, ABT199 (**B**), or the MCL-1 inhibitor, S63845 (**C**). Blue indicates synergy. Red indicates antagonism. (**D** and **E**) Relative cell viability, as measured using CellTiter-Glo, of the indicated isogenic cell lines that were treated with the indicated concentrations of ABT199 for 5-7 days.

*Supplementary Figure S5*

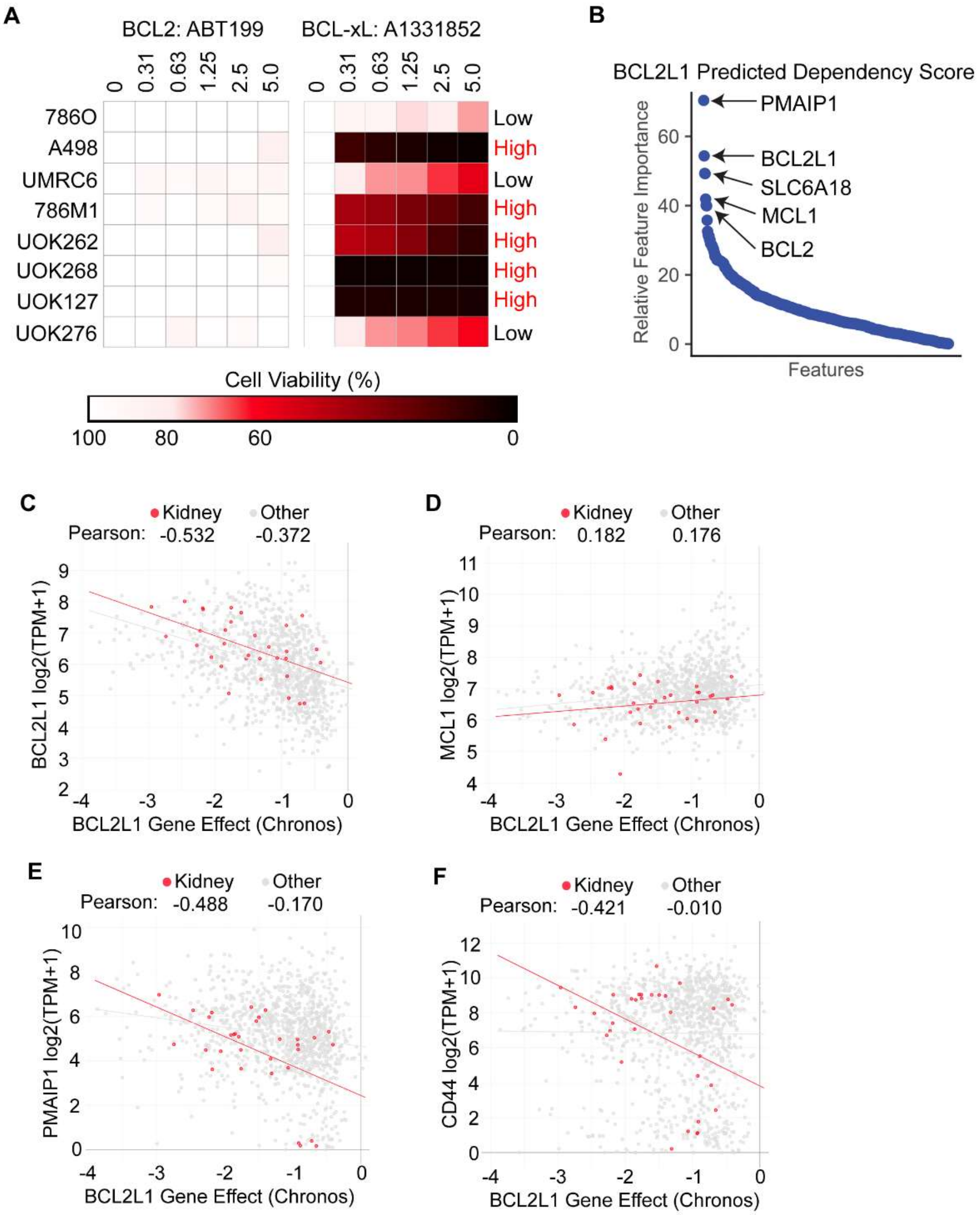


**Supplementary Figure S5. Gene Correlations Derived from the DepMap Datasets.** (**A**) Heatmap of the relative cell viability across the indicated cell lines treated with serial dilutions of either ABT199 (left) or A1331852 (right). Concentrations are expressed in μM. (**B**) Relative weight of the indicated genes in the correlation analysis using the DepMap dataset for the top predicted gene expression changes that are associated with BCL-$X_L$

dependence (note that a negative score means higher expression is associated with a more negative Chronos/Demeter score – greater BCL-$X_L$ dependence). Hockey stick plot computing the relative weight of correlates, including several members of the BCL-2 family. (**C** to **F**) Pearson correlation coefficient measured using the DepMap dataset for *BCL2L1* (**C**), *MCL1* (**D**), *PMAIP1*/NOXA (**E**), and *CD44* (**F**) expression versus BCL-$X_L$ dependence (Chronos scores). Red trendlines indicate the correlation of the indicated genes in the kidney lineage cells, compared to the lineage agnostic trendlines plotted in gray.

*Supplementary Figure S6*

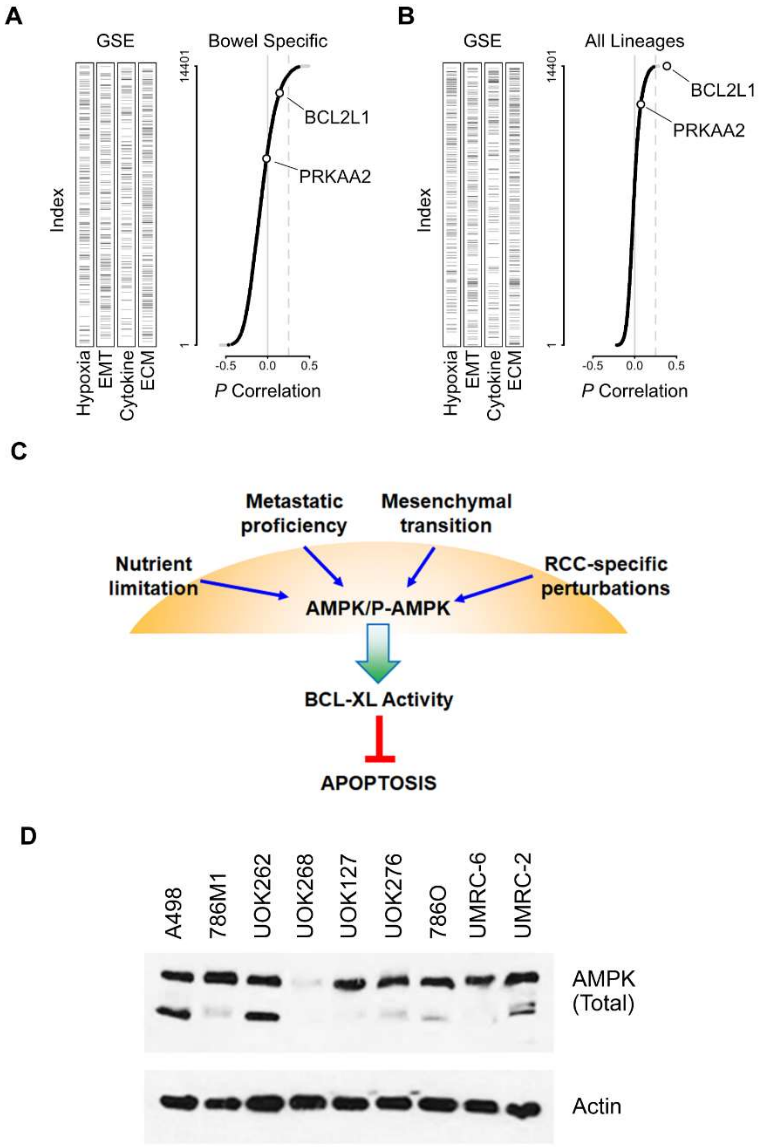


**Supplementary Figure S6. AMPKα2 Activation is a Kidney Lineage Biomarker of BCL-X$_L$ Dependence.** (**A**) Pairwise Pearson correlation values between bowel lineage-specific gene expression and *BCL2L1* dependence (right) and ranked enrichment of the indicated pathways (left; EMT, epithelial-to-mesenchymal transition; ECM, extracellular matrix). Data points in black are correlations which scored as kidney-specific compared to 19 other lineages and 20 quasi-lineages. Correlations were multiplied by -1. (**B**)

Lineage-agnostic Pearson correlations as described in (**A**). (**C**) Schematic of pathways relevant to *BCL2L1* dependence in kidney cancers and their convergence on AMPK activity, based on literature surveys. (**D**) Immunoblot analysis of the indicated proteins in the indicated renal cancer cell lines.
